# Photoperiod regulates gonadotrope cell division in medaka via melatonin, Tsh and folliculostellate cells

**DOI:** 10.1101/2023.06.09.544159

**Authors:** Muhammad Rahmad Royan, Kjetil Hodne, Rasoul Nourizadeh-lillabadi, Finn-Arne Weltzien, Christiaan V. Henkel, Romain Fontaine

**Affiliations:** Department of Preclinical Science and Pathology, Faculty of Veterinary Medicine, Norwegian University of Life Sciences, 1433 Ås, Norway

**Keywords:** photoperiod, seasonal reproduction, Lh, Fsh, Tsh, gonadotropes, plasticity, melatonin, folliculostellate cells, proliferation

## Abstract

In vertebrates, pituitary gonadotropins (follicle-stimulating and luteinizing hormones: FSH and LH) regulate gonadal development and maturation, therefore playing an essential role in reproduction. The seasonal regulation of gonadotropins has been widely studied in mammals and birds, and in these taxa thyroid-stimulating hormone (TSH) was found to play a critical role. By contrast, the seasonal regulation of gonadotropins remains unclear in teleost fish. In addition, the seasonal regulation of gonadotrope (gonadotropin-producing cell) proliferation has not been elucidated in any vertebrate group. Using the teleost fish medaka as a model, we show for the first time that long photoperiod enables reproduction by stimulating gonadotropin mRNA synthesis and gonadotrope cell proliferation. In female medaka, this proliferation is achieved by gonadotrope mitosis. We then demonstrate that in female medaka, photoperiod stimulates gonadotropin mRNA production and mitosis via an indirect intra-pituitary pathway, involving pituitary Tsh cells. We show that non-endocrine folliculostellate cells in the pituitary mediate the Tsh signal regulating gonadotrope activity and proliferation, as they are the only pituitary cells to express Tsh receptors and send projections to gonadotropes. Finally, we show that melatonin suppresses pituitary *tshba* expression in fish exposed to long photoperiod, suggesting that short photoperiod inhibits gonadotropin synthesis via melatonin in both fish and mammals. This study therefore demonstrates that in fish, photoperiod regulates gonadotrope cell activity and mitosis via a melatonin-Tsh pathway. It also reveals the existence of a novel intra-pituitary pathway for seasonal regulation of gonadotropes, involving folliculostellate cells, which we propose might also exist in other vertebrates.

**SIGNIFICANCE:** In seasonally breeding mammals and birds, the production of the hormones that regulate reproduction (gonadotropins) by gonadotropes is controlled by the pituitary thyroid-stimulating hormone (TSH) through an indirect pathway via the brain. However, in fish, how seasonal environmental signals influence gonadotropins remains unclear. Here, we show that in a long day seasonally breeding fish, medaka, photoperiod not only regulates the activity (hormone production) of the gonadotropes but also their proliferation. We also reveal a novel intra-pituitary pathway that regulates gonadotrope cell activity and number. This pathway involves melatonin, Tsh, and folliculostellate cells. Interestingly, as all these components are also found in the mammalian pituitary, this study suggests the existence of an alternative regulatory mechanism of seasonal gonadotropin production across vertebrates.

## INTRODUCTION

Reproduction is the most important biological function for all living organisms. For many animals, synchronization of reproduction with favorable environmental conditions is essential to ensure food abundance and moderate climate when their offspring are born in spring/summer season (1). For instance, animals with a long gestation (sheep and goat) breed between early autumn and winter, so that the offspring are born in spring or summer when grasses are abundant (2). In fish, synchronization of reproduction with favorable environmental conditions is also required. For instance, Atlantic salmon reproduce in the fall with their offspring hatching in spring when water temperatures are higher and food availability starts to increase (3).

In vertebrates, all physiological events associated with reproductive function are regulated by the brain-pituitary-gonad axis. In the brain, the hypothalamic gonadotropin-releasing hormone (GnRH) neurons integrate environmental and internal cues, and stimulate the synthesis and release of two gonadotropins (Follicle-stimulating hormone, FSH; and luteinizing hormone, LH) produced by gonadotropes in the pituitary (4). Gonadotropins are then secreted into the bloodstream to stimulate gametogenesis and steroidogenesis in the gonads (5). In addition to the hypothalamic control, gonadotropin production and release are also regulated by feedback signals from gonadal sex steroids, which act either directly on the gonadotropes or indirectly via the brain (e.g. the GnRH neurons; (6)). This system is highly conserved in vertebrates, except for a few differences at high taxonomic levels. For example, in fish, but not in mammals, Gnrh neurons directly innervate the anterior pituitary (7), and Fsh and Lh are produced by two distinct gonadotrope cell types (8, 9). These characteristics make teleost fish ideal models to investigate the differential regulation of the two gonadotropin hormones and gonadotrope plasticity and development.

One of the seasonal cues, daylength (photoperiod), is considered to be a noise-free (an almost identical and predictable rhythm each year) and primary signal for the activation of the reproductive axis in most seasonal breeders (10, 11). In mammals, photoperiodic information received by the pineal gland in the brain fine-tunes the secretion of melatonin hormone, whose duration and amplitude regulate thyroid-stimulating hormone (TSH) production in the pituitary *pars tuberalis* (PT; (12)). This PT-TSH signal then regulates gonadotropin production in the pituitary *pars distalis* (PD) via a retrograde pathway in the brain involving hypothalamic Kisspeptin, RFRP3, and GnRH neurons (12–15). In birds, the same pathway exists, except that PT-TSH is regulated by deep photoreceptors that directly sense photoperiod instead of melatonin (12, 13).

Although photoperiod regulates reproduction and gonadotropin levels in several fish species (e.g. stinging catfish (16), damselfish(17), and honmoroko (18)), little is known about the pathways involved. Teleost fishes do not possess an anatomically distinct PT (11), but some have a structure below their hypothalamus called the *saccus vasculosus* (SV) (19). The SV is proposed to process photoperiodic signals and to produce TSH in masu salmon (13, 20), although its involvement in reproduction remains unclear. In addition, nothing is known about the pathway regulating gonadotropes in teleost species that do not possess the SV (19). In Atlantic salmon, the pituitary product of the *tshbb* gene has been proposed to be comparable to PT-TSH in mammals and birds (21). *tshbb* is regulated by photoperiod (22), and has orthologs in many non-Salmonidae species, including zebrafish and medaka (23). In addition, several melatonin receptor isoforms show photoperiod-dependent expression levels in the pituitary in several teleost species, including in medaka (24, 25) and Atlantic salmon (26). Nonetheless, evidence for the functional role of pituitary Tsh and melatonin receptors in teleost reproduction remains scant.

Finally, despite the photoperiodic effect on *fshb* and *lhb* levels, it is not clear whether this regulation is due to increased gonadotrope cell activity, cell number, or both. Indeed, previous studies suggested that changes in Fsh and Lh cell number sometimes occur in the fish pituitary, allowing a rapid increase of *fshb* and *lhb* levels when increased activity of existing cells is not sufficient (27–29). While changes in pituitary gonadotrope cell number are usually explained by mitosis, transdifferentiation, differentiation of progenitor cells, and apoptosis (6, 30), how the seasonal photoperiod signal controls gonadotrope cell number is not known.

In this study, we used Japanese medaka (*Oryzias latipes*), a long-day seasonal breeder, as a model to investigate how photoperiod affects reproductive functions and gonadotrope cell plasticity. The species is a powerful model for genetic and developmental studies (31–33) due to a wide range of resources, as well as genetic and molecular techniques. These include, for instance, the recently developed 3D atlas of the pituitary, which facilitates visualization of all endocrine cell populations (29), and transgenic lines in which endogenous *lhb* and *fshb* promoters control the synthesis of fluorescent reporter proteins (34, 35). Here, we take advantage of these tools to elucidate how photoperiod regulates gonadotrope proliferation.

## RESULTS

### Long photoperiod stimulates reproduction and increases gonadotrope cell number

To evaluate the effect of photoperiod on reproductive capacity and gonadotrope cell number, we exposed adult fish to two different light regimes: Short (10 hours of light, SP) and long (14 hours of light, LP) photoperiods. After 3 months in these conditions, we counted the percentage of spawning females, shown by oviposited eggs, over seven days. While females in SP did not spawn at all, between 30% and 50% of the females in LP spawned each day (Supp. Fig. 1B-C), confirming that LP stimulates reproduction in medaka.

**Figure 1.**
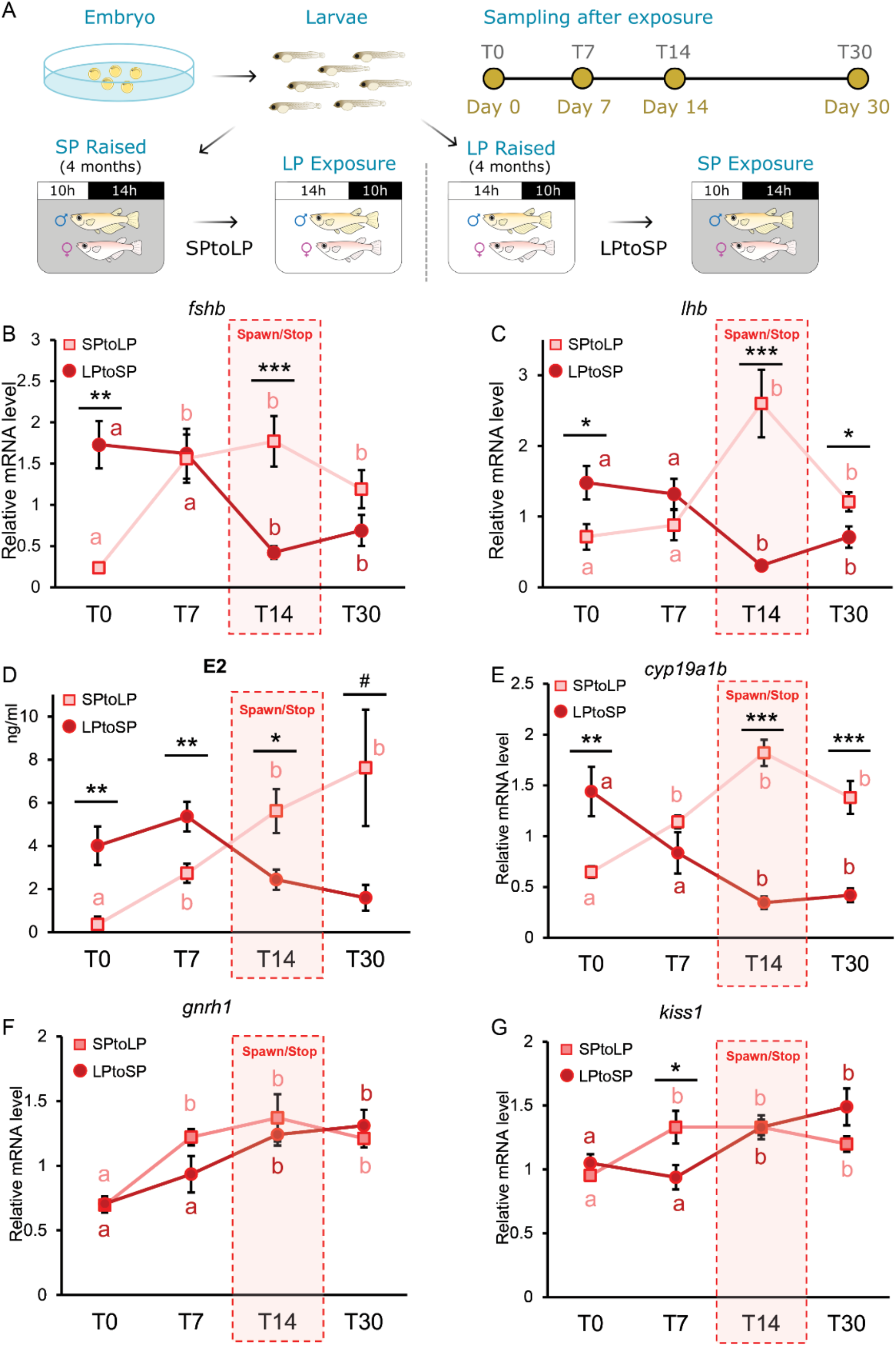
Long photoperiod induces an increase in gonadotrope cell activity and circulating sex steroids. (A) Illustration showing the experimental design. Fish were raised from hatching in LP or SP for 4 months before the photoperiod was changed. Fish were sampled on day 0 (just before the photoperiod change), 7, 14, and 30. (B-D) The fluctuation of *fshb* and *lhb* expression as well as E2 levels observed in females at different time points following the photoperiod change (n = 6-11). (E-G) The fluctuation of *cyp19a1b* levels in the pituitary as well as *gnrh1* and *kiss1* in the brain at different time points following the photoperiod change (n = 4-9) Black asterisks display statistically differences between the SPtoLP group and the SPtoLP group for each observation day. The letters (a, b) show statistically differences between each time points and T0 which is taken as reference within one treatment group. The statistical analyses were performed using two-sample independent t-test or Mann-Whitney U test (* < 0.05; ** < 0.01; *** < 0.001).

Consistent with the reproductive activity observed in fish kept in LP condition, both males and females kept in LP showed a significantly higher gonadosomatic index (GSI) compared to those kept in SP (Supp. Fig. 1D). This suggests that LP stimulates gonadal development, which is in line with the higher *fshb* and *lhb* levels observed in LP fish compared to SP fish (Supp. Fig. 1E-F). In addition, the pituitaries from LP fish contain a significantly higher number of Fsh and Lh cells than pituitaries from SP fish (Supp. Fig. 1G). This indicates that LP not only stimulates gonadotropin mNRA production, but also gonadotrope cell proliferation, and suggests that higher hormone-encoding mRNA production observed in LP exposed fish is at least partly due to gonadotrope hyperplasia. Furthermore, the percentage of Fsh and Lh cells in the pituitary of LP fish is the double that of SP fish (Supp. Fig. 1H) which supports their importance in reproduction under LP conditions.

We found that SP females did not reproduce at all when coupled with LP males. This interesting finding confirms that photoperiod length strongly regulates reproduction in female medaka (Supp. Fig. 1I). In males however, despite the lower levels of *fshb* and *lhb*, number of gonadotropes, and GSI, SP males were still able to reproduce, as shown by 7 out of 10 couples of SP males and LP females reproducing with 10-100% fertilized eggs. These results suggest that males are less affected by photoperiod for the control of their reproductive activity. For this reason, we focused on females in the rest of the study, and males are analysed separately in Supp. Fig. 2.

**Figure 2.**
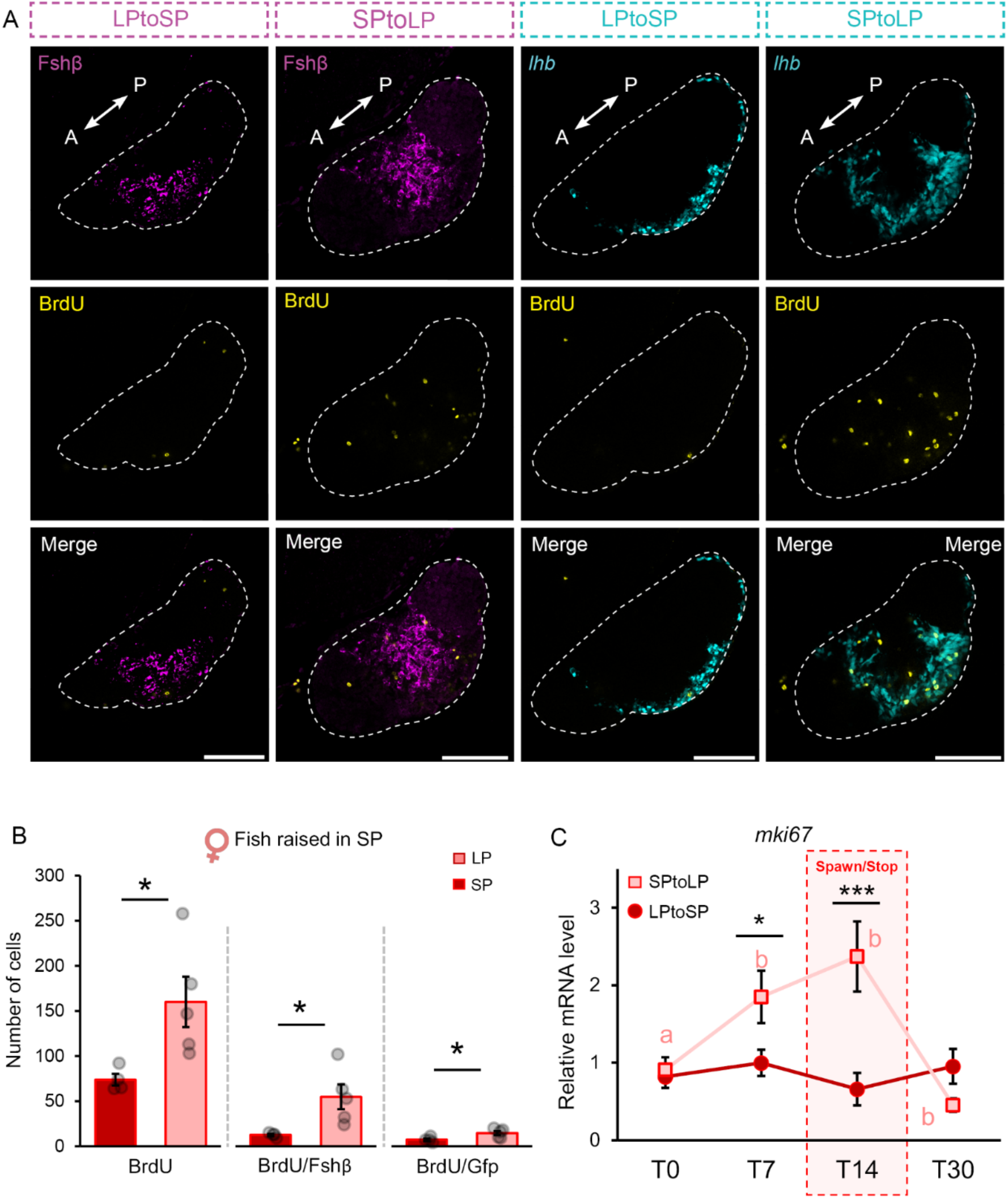
Long photoperiod induces an increase in gonadotrope cell mitosis in SP females. (A) Confocal planes illustrating Fsh (magenta) and Lh (cyan) cell mitosis using BrdU marker (yellow) 5 days after photoperiod change (LPtoSP and SPtoLP). Scale bar: 100 µm. The dashed line indicates the whole parasagittal plane of the pituitary (A: anterior, P: posterior). (B) Number of mitotic cells in SP females after exposing them to LP for 5 days or kept in SP for control (n = 4-5). (C) Relative expression levels of a mitotic cell marker, *mki67*, in females at different time points following the photoperiod change as described in Fig. 1A (n = 6-11). The statistical analyses were performed using two-sample independent t-test or Mann-Whitney U test, in which the graph represents mean ± SEM while the jittered dots represent each individual (* < 0.05; ** < 0.01; *** < 0.001).

### Changes in photoperiod induce rapid physiological modifications

To determine how quickly the physiological modifications occur following a change in photoperiod, we raised fish either in SP or LP for 4 months before switching them to LP or SP, respectively (Fig. 1A). After 14 days, SP raised fish exposed to LP (SPtoLP) and LP raised fish exposed to SP (LPtoSP) had started and stopped reproducing, respectively.

We then looked at gonadotropin mRNA levels (*lhb* and *fshb*) and observed that both decrease in LPtoSP females, but increase in SPtoLP females (Fig. 1B-C). While *fshb* levels are already significantly higher compared to day 0 in SPtoLP females after 7 days, significant changes on *lhb* levels are only observed after 14 days. In LPtoSP females, both *lhb* and *fshb* significant declined by day 14. These major changes in gonadotropin levels after 14 days likely explain why the fish started or stopped reproducing at this time point. These changes are generally consistent with the observed changes in estradiol (E2) levels which significantly increased in 7 days in SPtoLP females and trended down in LPtoSP females (Fig. 1D).

Meanwhile, the levels of *cyp19a1b* (encoding aromatase, which is known to be expressed in medaka pituitary gonadotropes (28) and folliculostellate cells (36) show the same pattern as the gonadotropins (Fig. 1E).

In the brain, *gnrh1* (which encode the primary stimulatory factors for Lh gonadotropes (37)) and *kiss1* levels, also significantly increase in SPtoLP females (Fig. 1F,G). These results suggest that a brain signalling pathway, similar to what has been described in mammals and birds, is also activated in medaka. However, both *gnrh1* and *kiss1* levels also significantly increase in LPtoSP fish, suggesting that the biological response to photoperiod must be coordinated downstream from the brain (e.g. in the pituitary).

### Long photoperiod stimulates gonadotrope proliferation via mitosis in females

Because the changes in *lhb* and *fshb* mRNA levels were found to significantly increase after increasing photoperiod from SP to LP, we investigated whether LP induced gonadotrope cell mitosis. We thus performed BrdU incorporation experiments to determine whether gonadotropes were proliferating (Fig. 2A). While we saw no change in mitotic cell number in the pituitary of LPtoSP fish 5 days after the light regime change, with a total of about 60 proliferative cell per pituitary (Supp. Fig. 3), we found that the total number of mitotic cells, as well as the number of mitotic Fsh and Lh cells significantly increased in SPtoLP females (Fig. 2B). This agrees with the increased mRNA levels of the proliferation cell marker *mki67* observed in females at day 7 and day 14 (Fig. 2C), clearly indicating that LP stimulates gonadotrope cell proliferation in females.

**Figure 3.**
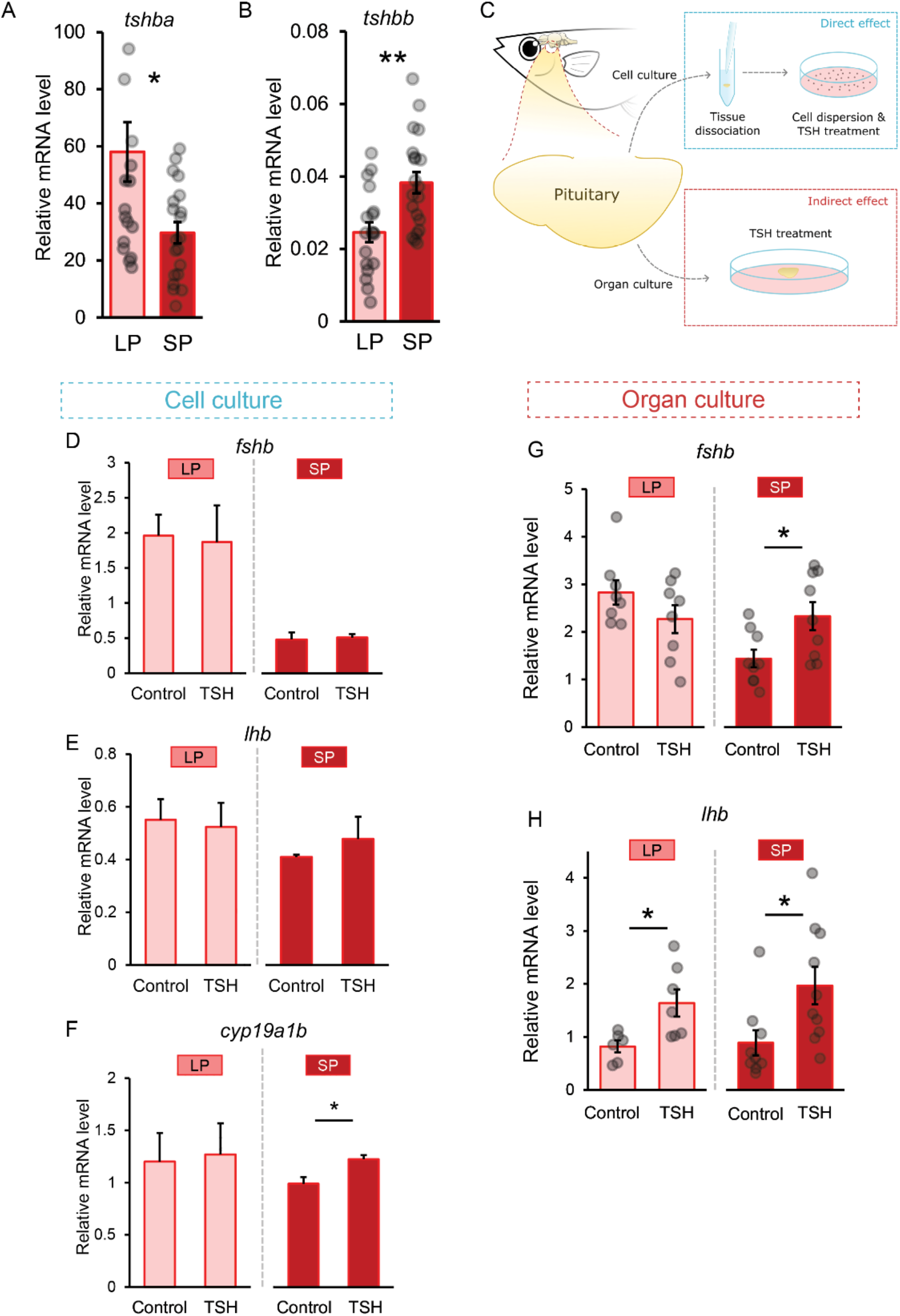
Tsh indirectly stimulates gonadotrope cell activity in SP females. (A-B) *tshba* (A) and *tshbb* (B) mRNA levels in LP and SP condition in female medaka (n = 19-20). (C) Experimental scheme illustrating the treatment of 0.5 µM of bovine TSH pituitary extract to dissociated cell culture and organ culture of medaka pituitary to evaluate direct and indirect effect of Tsh on gonadotrope cell activity. (D-F) Effect of TSH or vehicle (control) on *fshb* and *lhb* mRNA levels in dispersed pituitary cell cultures in female medaka from LP and SP condition (n = 3; in which each replicate represents 4 pooled pituitaries), the bar represents mean + SEM. (G-H) Effect of TSH or vehicle (control) on *fshb* and *lhb* mRNA levels in *ex vivo* medaka pituitary organ cultures of female medaka from LP and SP condition (n = 6-10). The statistical analyses were performed using two-sample independent t-test or Mann Whitney U test. All graphs (unless otherwise stated) are represented as mean ± SEM with the jitter dots representing each individual (* < 0.05; ** < 0.01; *** < 0.001).

### Tsh indirectly regulates gonadotrope cell activity

Since in mammals and birds the role of PT-TSH is crucial for the seasonal production of gonadotropins, we investigated whether Tsh also plays a role in regulating gonadotropin levels in medaka. Fish possess two paralog genes encoding proteins with similarity to TSHβ: *tshba* and *tshbb* (23), which both display photoperiod-dependent mRNA levels in females (Fig. 3A-B). While *tshba* levels were significantly higher in LP females than in SP females, *tshbb* levels were significantly lower (Fig. 3A,B). These results therefore are consistent with a role of *tshba* and *tshbb* in the photoperiod signalling pathway in fish. Of note, the levels for *tshbb* were much lower than those of *tshba* as shown by the Cq values (average Cq values *tshba* = 19; *tshbb* = 32) and in the medaka pituitary RNA-seq data where *tshbb* was not detected while high expression of *tshba* was found (Supp. Fig. 4A). While these results suggest that *tshbb* might have a more minor role than *tshba*, the fact that mammalian TSHβ has a closer identity to medaka Tshβa than Tshβb (Supp. Fig 4B-D), prompted us to investigate further the effects of Tsh on gonadotropin mRNA synthesis.

**Figure 4.**
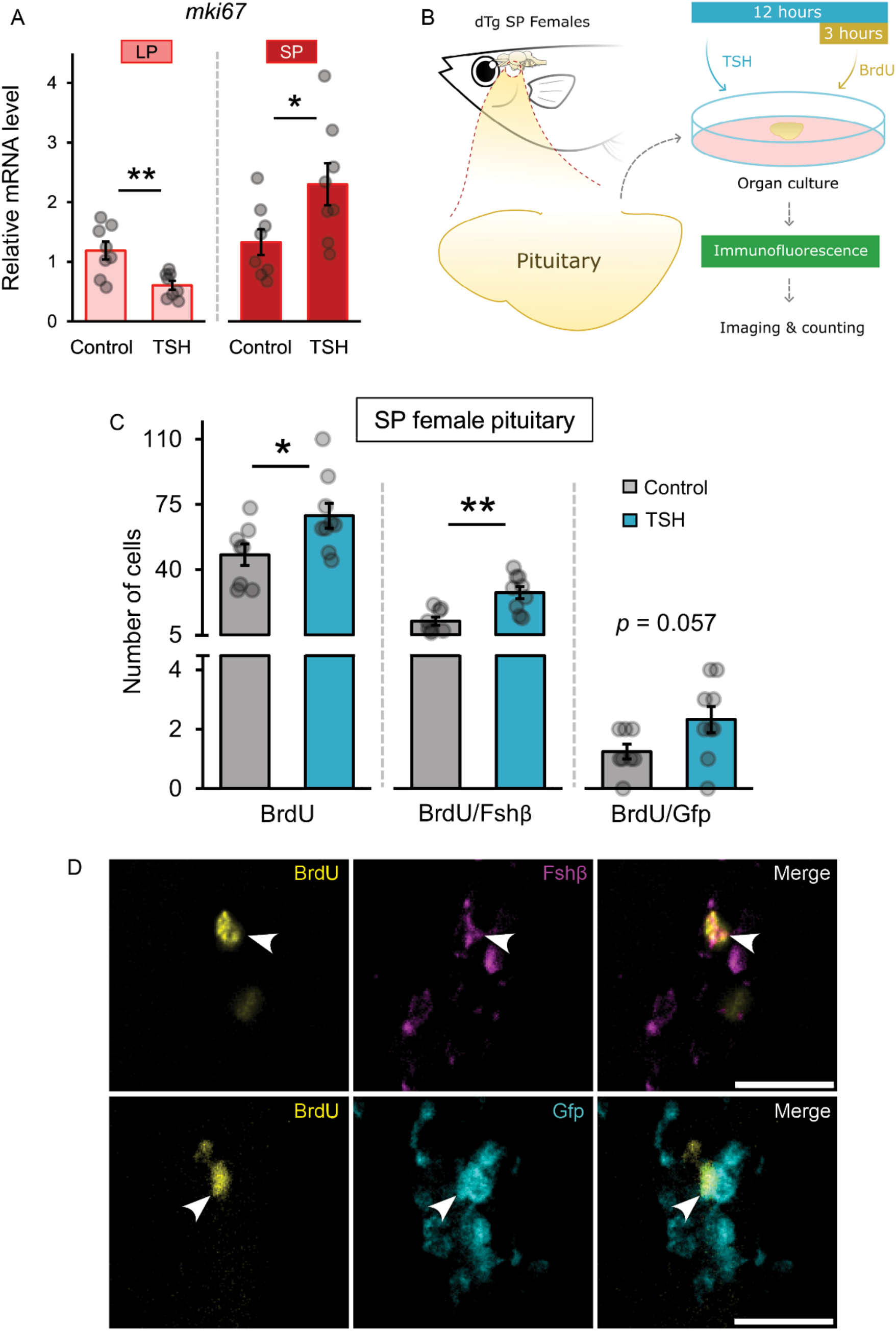
Tsh stimulate mitotic activity in the pituitary of SP females. (A) Effect of 0.5 µM of bovine TSH pituitary extract on *mki67* levels in the female medaka pituitary organ cultures (n = 6-10) in LP and SP condition. (B) Schematic illustration of *ex vivo* medaka pituitary organ culture with bovine TSH pituitary extract and BrdU treatment to evaluate gonadotrope mitosis. (C) Number of mitotic cells in SP females exposed to TSH for 12 hours (n = 8-9). (D) Visual representation from the pituitary of dTg fish with Gfp signal for Lh cell (cyan), Fshβ for Fsh cells (magenta), and BrdU for mitotic cells (yellow) in the pituitary of SP females treated with TSH, as taken by Thunder fluorescence microcsope. The statistical analyses were performed using two-sample independent t-test or Mann Whitney U test, in which the error bar represents SEM while the jittered dots represent each individual (* < 0.05; ** < 0.01). Arrows point the mitotic cells. Scale bar: 20 µm.

We applied bovine TSH pituitary extract (bovine) to a dispersed pituitary cell culture (*in vitro*) in order to determine the direct effects on gonadotropes, and to pituitary organ cultures (*ex vivo*) to identify indirect effects (Fig. 3D-H). In dispersed pituitary cell cultures from females, we did not observe any change in *fshb* and *lhb* levels in any condition after TSH treatment (Fig. 3D,E). However, TSH stimulated *cyp19a1b* mRNA levels in cells from SP females (Fig. 3F), suggesting that some aromatase-expressing pituitary cells are directly stimulated by TSH. Meanwhile, in pituitary organ culture from females kept in SP condition, TSH significantly upregulated *fshb* and *lhb* levels (Fig. 3G,H) demonstrating that in females, Tsh regulates gonadotrope cell activity in the pituitary, indirectly through aromatase-expressing cells.

### Tsh stimulates gonadotrope mitosis in females

Because long photoperiod stimulates both gonadotrope cell division and activity in females, we then investigated whether Tsh regulates gonadotrope proliferation in pituitary organ cultures. First, TSH downregulated *mki67* levels in LP but upregulated levels in SP females (Fig. 4A). Next, we used BrdU treatment using pituitary from SP females (as shown in Fig. 4B) and found that both the total number of mitotic cells and the number of mitotic Fsh cells increase after 12-hour TSH treatment (Fig. 4C,D). TSH treatment also elevated Lh cell mitosis close to statistical significance (*p* = 0.057). These results suggest that Tsh stimulates both gonadotrope cell activity and their mitosis in females.

### Folliculostellate cells mediate Tsh regulation of gonadotrope cells

Considering the indirect regulation of Tsh on gonadotrope cell activity and proliferation, we decided to investigate which pituitary cells might mediate the Tsh signals to gonadotrope cells.

Using the available pituitary bulk transcriptomic data (which have higher sequencing depth than single cell sequencing), we found that of the two Tsh receptors identified in the medaka genome (23), only one is expressed above background level in the medaka pituitary (Supp. Fig. 5A). This receptor does not show photoperiod-dependent expression levels (Supp. Fig. 5B). Single cell pituitary transcriptomic data indicates that this Tsh receptor is specifically expressed in a small cluster of cells (Fig. 5A,B). This cell cluster also expresses common markers for non-endocrine folliculostellate (FS) cells (Supp. Fig. 6), including high levels of aromatase (Fig. 5C). Using RNAscope, we localized Tsh-receptor expression in the dorsal part of the pituitary. Tsh-receptor expression labelling completely coincides with the labelling of cells expressing high levels of aromatase (Fig. 5D-F), thus confirming the single cell transcriptomic data. It also agrees with the increased of aromatase mRNA levels observed following TSH stimulation in dispersed pituitary cell cultures from females kept in SP condition (Fig. 3F). All these results thus support the direct regulation of FS cells by Tsh cells, suggesting that FS cells mediate the regulation of gonadotrope cell activity and mitosis.

**Figure 5.**
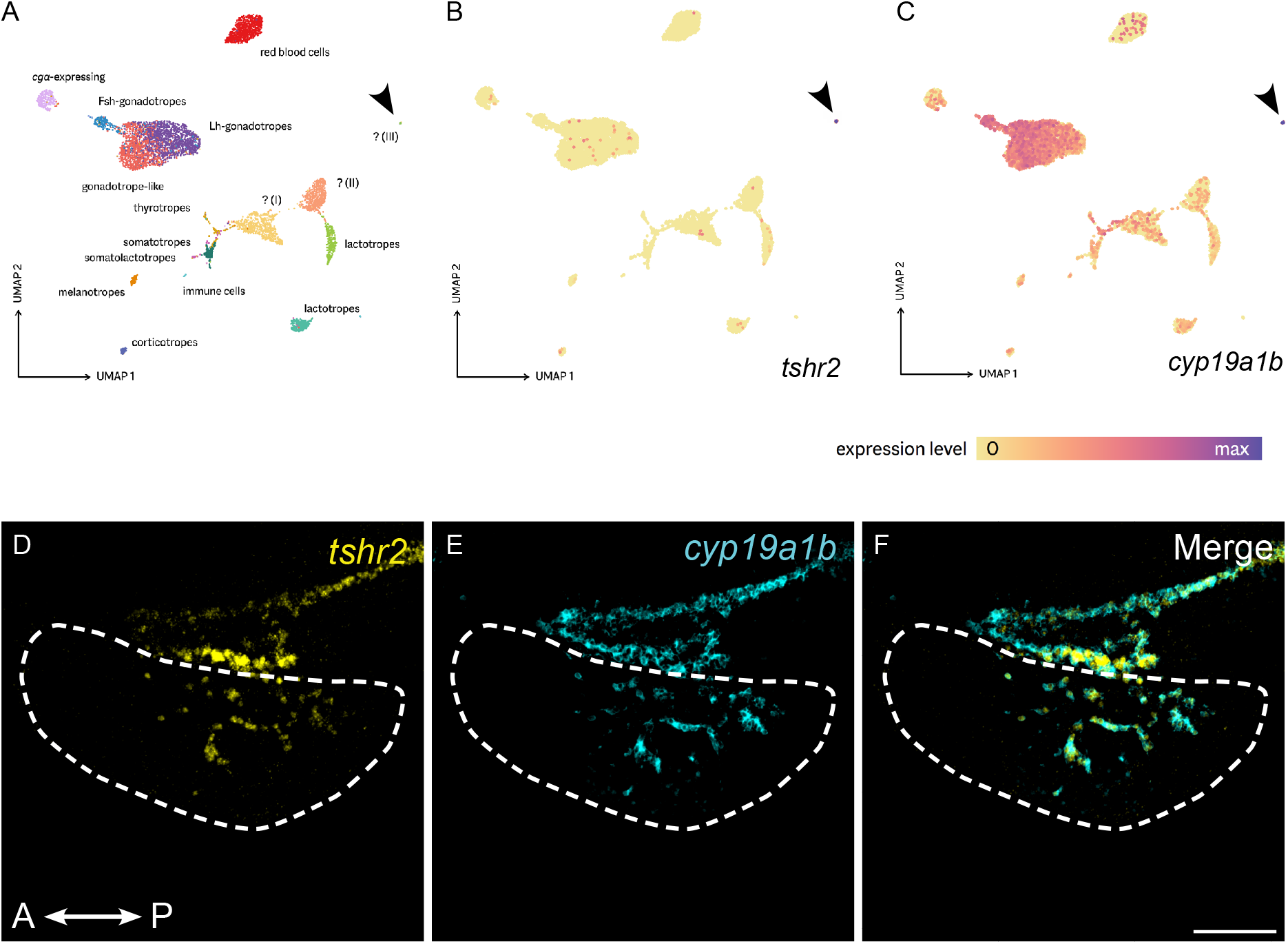
Tsh regulation of gonadotrope cells is mediated via folliculostellate cells. (A) The pituitary cell clusters identified by single-cell sequencing, showing three uncharacterized cell types (I-III). (B-C) *tshr2* and *cyp19a1b* are specifically expressed in one of these uncharacterized cell types (III), with *cyp19a1b* also showing modest expression in other pituitary cell types (e.g. gonadotropes). Figures are UMAP projections generated using Seurat. Colour scale: log-transformed relative expression level (maximized per gene). (D-F) Confocal planes showing co-expression of *tshr2* and *cyp19a1b* in the same pituitary cells. Parasagittal section. Scale bar: 100 µm.

**Figure 6:**
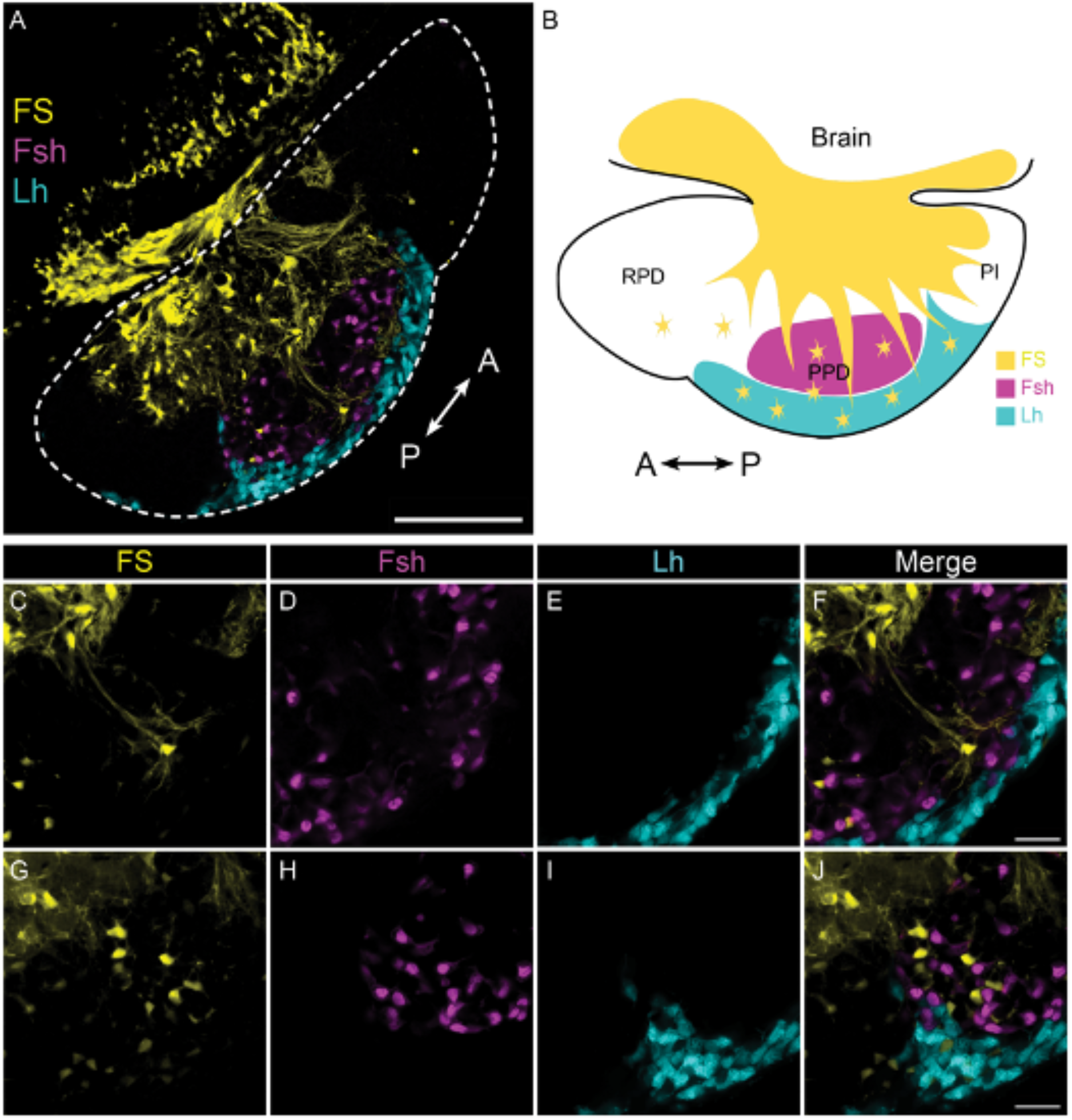
Folliculostellate cells connect to gonadotropes. Labeling of folliculostellate cells using the dipeptide β-Ala-Lys-Nε-AMCA in double transgenic (*lhb*-GfpII/*fshb*-DsRed2) medaka fish. (H) Schematic illustration of a parasagittal section of the medaka pituitary showing folliculostellate cell localization together with Fsh and Lh cells. (I-L) High magnification of confocal planes showing the projection of folliculostellate cell extensions (yellow) to gonadotrope cells (Lh: cyan; Fsh: magenta). (M-P) High magnification of confocal planes showing of folliculostellate cells that borders with gonadotropes. Dashed line indicates the whole parasagittal plane of the pituitary (A: anterior, P: posterior).

To further characterize the FS cells in the medaka pituitary, we labelled them using the commonly used dipeptide β-Ala-Lys-Nε-AMCA. FS cell bodies show a similar location distribution to the *cyp19a1b+/ tshr+* cells (Fig. 6A,B). We observed that these FS cells send projections to gonadotropes in the ventral adenohypophysis (Fig. 6C-F), and that some FS cells are in close proximity to gonadotropes (Fig. 6G-J). This organization is similar for females in both SP and LP conditions (data not shown). Together these results suggest that the Tsh signal is relayed to gonadotropes directly by FS cells.

### Melatonin suppresses Tsh cell activity in the medaka pituitary

We also investigated whether melatonin regulates Tsh production in medaka as it does in mammals. First, we confirmed previous results by showing that among the four melatonin receptors that have been described in medaka (24, 25) (Supp. Fig. 5C-F), two (*melr1a* and *melr1a*-*like*) show photoperiod-dependent expression levels, with higher levels in SP fish compared to LP fish (although only *melr1a-like* significantly declined in the present study, Supp. Fig. 5D).

We then investigated whether melatonin regulates *tshba* or *tshbb* expression in dissociated pituitary cell culture or in pituitary organ culture (Fig. 7). In cell culture from female pituitaries, we did not observe any direct effect of melatonin treatment on either *tshba* or *tshbb* levels (Fig. 7A,B). In pituitary organ culture however, in agreement with a previous study where melatonin inhibited *tshba* levels in females (38), we found that melatonin significantly reduced *tshba* levels in males (Supp. Fig. 2) and tended to decrease levels in females (p = 0,093) (Fig. 7C). Contrary to *tshba* levels, *tshbb* expression levels were significantly upregulated by melatonin in pituitary organ cultures from LP females (Fig. 7D). Together, these results support a role of pituitary melatonin receptors in photoperiod signal integration in the pituitary through the regulation of Tsh synthesis and on gonadotrope cell proliferation.

**Figure 7.**
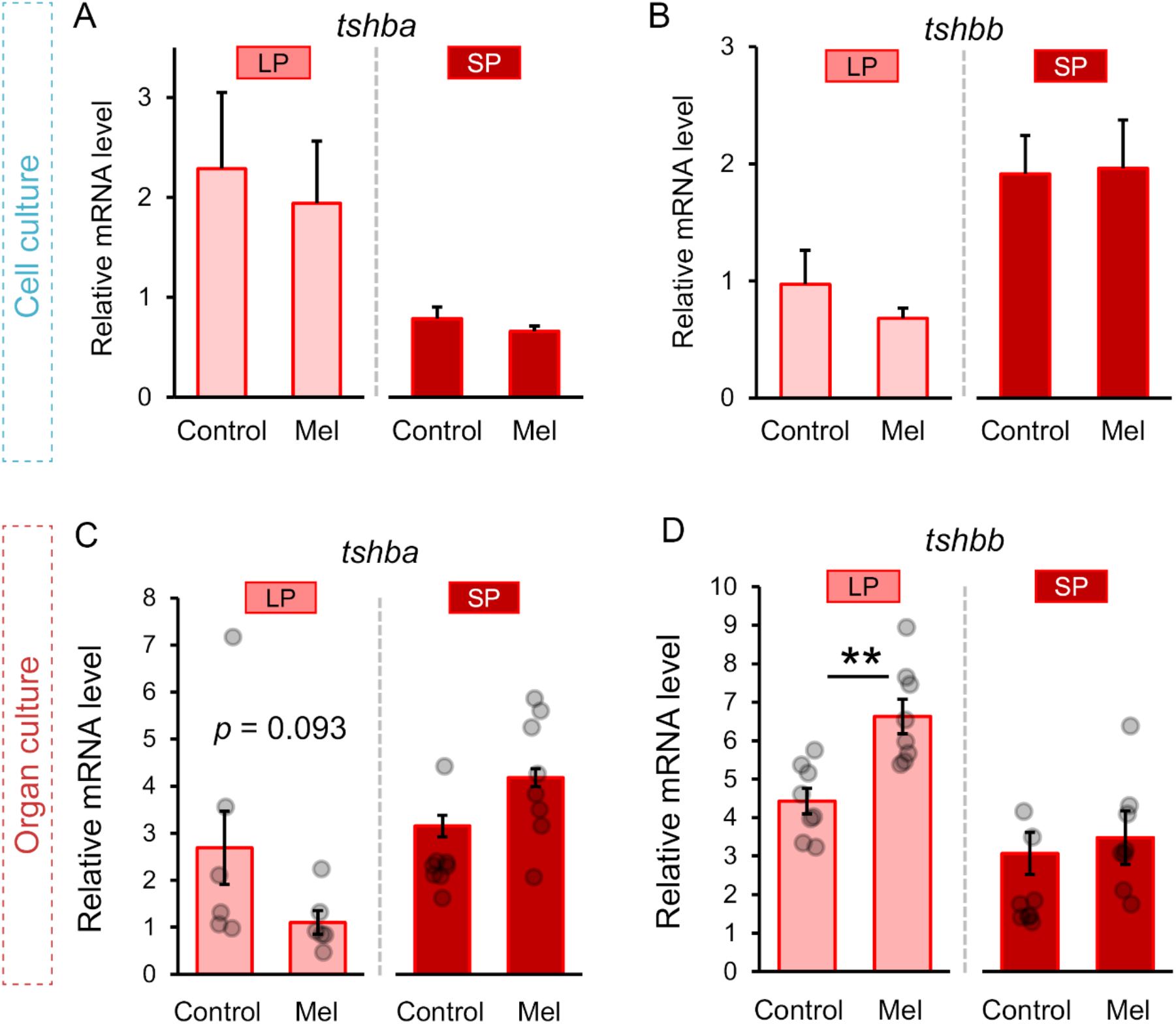
Melatonin indirectly regulates Tsh cell activity. (A-B) Effect of 10 µM melatonin on *tshba* and *tshbb* levels in dispersed cell cultures of medaka pituitaries from LP and SP condition (n = 3 cultures, each n represents 4 pooled pituitaries). (C-D) Effect of 10 µM melatonin on *tshba* and *tshbb* levels in medaka pituitary organ cultures from LP and SP condition (n = 6-10). The statistical analyses were performed using two-sample independent t-test or Mann Whitney U test, in which the error bar represents SEM while the jittered dots represent each individual (* < 0.05; ** < 0.01).

## DISCUSSION

While seasonal regulation of gonadotropin production in mammals and birds has been widely studied (12, 13), this process is still poorly understood in fish. Using the teleost model organism the Japanese medaka, a species which reproduces in the wild during the summer when photoperiod is long and temperatures are warm (39), we investigated how photoperiod regulates gonadotropin mRNA production and gonadotrope cell number, as well as the regulatory pathways involved.

Following the confirmation in our laboratory that, as previously shown by (39, 40), medaka reproduction is induced by an increase in photoperiod length, we found that gonadal development is stimulated under LP conditions via increased gonadotropin mNRA production in the pituitary. This agrees with observations in other teleost species (41–43), as well as in mammals (44). In addition, we show here that the increase in gonadotropin transcripts is at least partly due to an increase of the gonadotrope cell number. In previous studies, we have indeed shown that changes in hormone production are often in line with changes in cell number (27-29, 45). Nevertheless, this is to the best of our knowledge, the first report of seasonal regulation of gonadotrope hyperplasia by photoperiod in any vertebrate.

Interestingly, while SP completely blocks reproduction in females, it does not in males, despite decreased *fshb* and *lhb* expression as well as lower Fsh and Lh cell numbers. Together with the fact that *fshb* or *fshr* knockout medaka halt vitellogenesis in the female ovary, resulting in infertility due to immature follicles (46, 47), our results support that gonadotropins play a pivotal role for ovarian development and maturation in females. However, in agreement with a previous study in which male medaka produced mature spermatozoa even under SP conditions (39), spermatogenesis still occurs in SP medaka males in our study, even though fewer gametes or spermatozoa might be produced in those males, as indicated by the lower GSI compared to LP males. In addition, the fact that SP males are still able to reproduce suggests that spermatogenesis may not be completely dependent on gonadotropins, and that other factors could compensate for low gonadotropin levels. This hypothesis is consistent with previous knockout studies in male medaka, which demonstrated that the loss of *fshb* or *lhb* (46) or double knockout (KO) of *fshr* and *lhr* (47, 48) does not disrupt spermatogenesis. In male zebrafish, although testis development is delayed significantly, Fsh-deficient fish are still fertile (49). It was demonstrated that Lh could compensate for Fsh deficiency by activating the Fsh receptor in zebrafish (50, 51), suggesting that low levels of either or both Fsh and Lh hormones could still, to some extent, stimulate spermatogenesis. Meanwhile, double KO of *fshb* and *lhb* in zebrafish males causes infertility (51), which suggests that the essentiality of gonadotropins depends on the species. Interestingly, in an *in vitro* experiment, human FSHR-expressing cells could be stimulated not only by human-FSH, but also by human-TSH, in a dose-dependent manner (52). TSH was also shown to have a pronounced effect on the development of chicken testis (53). As *tshba* levels increase in males (but not in females) in SP conditions, where the males still produce sperm, we hypothesize that *tshba*-derived protein (called Tsh throughout the rest of this discussion) compensates for the low Fsh and Lh levels in SP, thus stimulating spermatogenesis.

Following the observation of gonadotrope hyperplasia in LP males and females, we have demonstrated for the first time that photoperiod can regulate gonadotrope cell mitosis in vertebrates. Although stimulation of gonadotrope proliferation by photoperiod has never been shown before in any vertebrate, mitosis of gonadotrope cells has previously been observed in the medaka pituitary. In fact, our previous studies revealed an important role of gonadal sex steroids in the regulation of gonadotrope mitosis in the medaka pituitary (28, 45). Surprisingly, despite increased Fsh and Lh cell numbers in LP males compared to SP males, we found that LP actually inhibited gonadotrope mitosis in males. The difference between females and males could simply result from a sexual dimorphism in the timing of proliferation, as gonadotrope cell mitosis was only investigated at one time point (five days after the change in photoperiod).

In ewes, it has been shown that cell division in the pituitary *pars tuberalis* decreased in short photoperiod, although the cell types were not identified (54). By contrast, photoperiod change does not affect cell division in the pituitary PT in male sheep (55), suggesting that sex differences in cell division by the effect of photoperiod also occur in other vertebrate species. However, it is also possible that LP-induced gonadotrope cell hyperplasia results from another mechanism in males. Indeed, we previously demonstrated that Fsh cell hyperplasia originates from Fsh cell mitosis in both sexes, as well as transdifferentiation of Tsh cells in females, while a decreasing Sox2-immunolabeled cell population in males suggested that progenitor cells may contribute in gonadotrope proliferation in males (45). Thus, it would not be surprising if males exposed to LP use another mechanism than mitosis, such as the differentiation of progenitor cells, to increase gonadotrope cell number.

In mammals and birds, *pars tuberalis* (PT)-TSH plays an essential role in regulating gonadotropin production indirectly via a retrograde pathway through the brain. Interestingly, *tshba* and *tshbb* levels are also photoperiod-dependent in medaka. The mRNA levels of *tshbb* are consistently upregulated in SP medaka. This is opposite to in Atlantic salmon, where *tshbb,* which is strongly associated with smoltification (21), is upregulated in LP conditions (22), suggesting species differences. For medaka as a non-smoltifying species, *tshbb* upregulation concurrently with *fshb* and *lhb* suppression in SP medaka suggest that *tshbb*-expressing cells (further called Tsh-like cells throughout) might play an inhibitory role for reproduction, at least in females. Very little is known about Tshβb, apart from that the protein sequence is quite different than Tshβa. For instance, we do not know if it binds to the known Tsh receptors or to other types of receptors. Nevertheless, the extremely low levels compared to all other pituitary hormones suggest that Tsh-like might be used as a paracrine signal linked to photoperiodism, and not an endocrine signal.

We found that *tshba* is expressed at a much higher level than *tshbb* in the medaka pituitary, and furthermore the effect of photoperiod on *tshba* expression is sexually dimorphic. In females, *tshba* levels are high in LP conditions, while they are constantly low in males. Previous studies have linked *tshba* and photoperiod in fish. For instance, LP was found to stimulate *tshba* in medaka (56), although the study did not specify the sex. Increase of *tshba* in medaka is also associated with mating experience in males (57), suggesting that Tsh cells may play a role in medaka reproduction. In chub mackerel, *tshba* levels are also upregulated in LP fish, and are linked to the increase of *fshb* and *lhb* levels in both sexes (58). Here, we show that in addition to be controlled by photoperiod, Tsh regulates gonadotropin mRNA production in medaka of both sexes, and gonadotrope mitosis in females.

Interestingly, in both males and females, the stimulatory effects of Tsh on gonadotropin mRNA levels were observed on *ex vivo* isolated pituitaries, but not in dissociated cells. Therefore, in addition to the effects observed in the brain, as expected according to other mammalian and bird studies, we demonstrate here for the first time the existence of an indirect, intra-pituitary pathway for the regulation of gonadotropin mRNA production by Tsh. We identified a subset of folliculostellate (FS) cells to be this intermediate, as these are the only pituitary cells expressing Tsh-receptors. Our single cell data clearly shows that what is regularly named FS cells according to the usual markers (e.g. *s100*), represents in fact an heterogenous cell population, as these are expressed in at least three distinct cell populations. As FS marker genes have been established based on bulk analysis of β-Ala-Lys-Nε-AMCA-stained cells, this complex presumably also includes pituicytes (59, 60). Only one small cell cluster in this complex expresses the Tsh receptor, as well as high levels of aromatase. Our study therefore provides a starting point for further characterizations and functional analyses of FS/pituicyte heterogeneity.

Expression of Tsh receptors has also been shown in FS cells from human (61, 62), mouse (63), and chicken pituitary (64), suggesting that this intra-pituitary pathway for Tsh regulation of gonadotrope cells, mediated by FS cells, might be conserved among vertebrates. FS cells possessing cytoplasmic extensions have been suggested to assist in paracrine communications among pituitary cells (65–68). The previously described gap junction-mediated calcium (Ca^2+^) propagation among FS cells (66, 69), as well as gap junction-mediated communication between FS and other endocrine cells (70, 71), may enable a signal transduction cascade that induces gonadotrope cell activity and mitosis.

Finally, we show that, like in mammals and in agreement with previous studies (72), pituitary melatonin receptors may play a role in seasonal photoperiodism in medaka as they are differentially regulated by photoperiod. This is also supported by our finding that melatonin downregulates *tshba*, as previously shown in females (38). This is intriguing as contrary to in females, SP increased *tshba* levels in males, suggesting that another factor, more powerful than melatonin, is regulating *tshba* levels in males. Finally, while the cellular identity of the cells expressing melatonin receptors remains to be determined in fish, the fact that this regulation is observed *in vivo* and in *ex vivo* pituitary cultures but not in *in vitro* dissociated pituitary cell cultures suggests that the effect of melatonin on thyrotropes is indirect. Also, deep photoreceptors have been found in all the tissues that have been investigated in fish, including the pituitary (73, 74), but whether they play a role in Tsh regulation as in birds remains to be investigated.

In summary, because gametogenesis still occurs in SP males the role of *tshba*, FS cells, and melatonin in the regulation of gonadotrope cell activity and proliferation is less clear in males than in females. In females, we propose the following hypothesis (Fig. 8): during winter conditions (short photoperiod), high levels of melatonin indirectly inhibit Tsh cells thus suppressing Tsh stimulation of gonadotropin production and gonadotrope cell proliferation, thereby blocking reproduction. Melatonin also stimulates Tsh-like cells for which the potential role in the control of gonadotropin production remains to be investigated. In summer conditions (long photoperiod), reduced melatonin levels permit the activation of the Tsh cells which stimulate gonadotrope cell activity and mitosis via FS cells. The increase of gonadotrope cell mitosis increases gonadotrope cell number, which contributes to the elevation of gonadotropin production necessary for gonadal development and maturation, and thus for the fish to reproduce.

**Figure 8.**
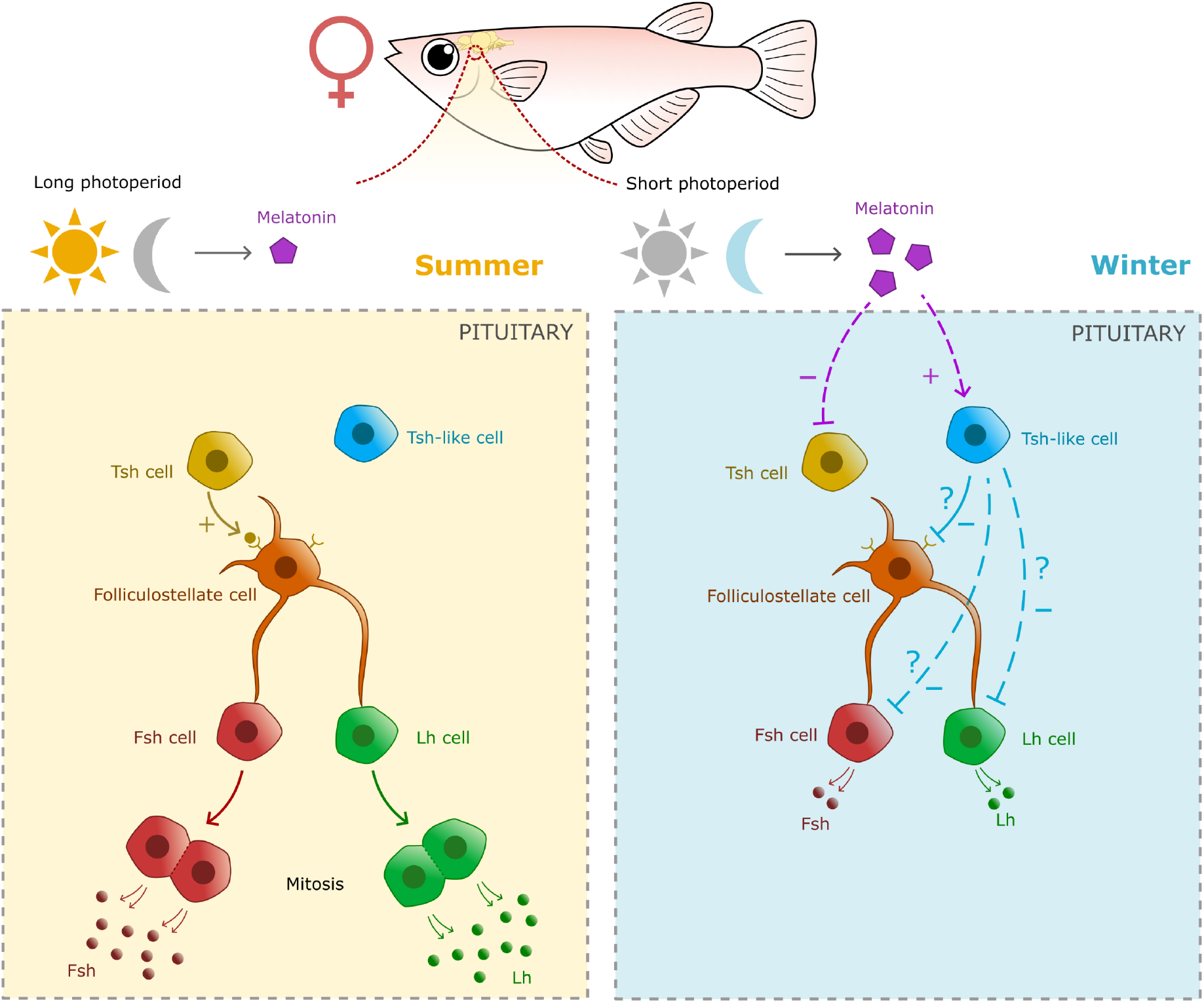
Schema of the proposed hypothesis on the photoperiodic regulation of gonadotrope cell mitosis via melatoinin, Tsh, and folliculostellate cells. In summer photoperiod (left panel), melatonin levels are suppressed, allowing Tsh cells to stimulate gonadotrope cell activity and mitosis, via folliculostellate cells. As a result, increasing number of gonadotrope cell due to mitosis participates in the increase gonadotropin production necessary for gametogenesis and steroidogenesis, which allow the fish to reproduce. In winter photoperiod condition (right panel), high melatonin levels indirectly suppress Tsh cell activity. In absence of Tsh stimulation, gonadotrope activity and mitosis remain low. By contrast, melatonin indirectly upregulates Tsh-like cell activity that might inhibit Fsh and Lh, but this needs to be confirmed.

## MATERIALS AND METHODS

### Experimental animals

Male and female of wild type (WT) and double transgenic (dTg) (*lhb*:hrGfpII/*fshb*:DsRed2) line (35) medaka (Hd-rR strain) were raised at 28 °C in a re-circulating water system (pH 7.5; 800 µS) with 14 h light and 10 h dark (long photoperiod, LP) or 10 h light and 14 h dark (short photoperiod, SP). The photoperiod regimes were based on what has been previously performed in medaka to mimic seasonal variation (75). Fish were fed three times daily using artemia (twice) and artificial feed (once). The sex was determined based on secondary sexual characteristics (76).

### Photoperiod exposure *in vivo*

*Experiment 1*: To compare gonadotrope cell activity and number between LP and SP fish, 2-month-old medaka were distributed into two groups of mixed sex. The first group was kept in LP while the second group was kept in SP for 3 months before sampling. The number of females showing oviposited eggs was used as a measure of the percentage of spawning animals. *Experiment 2*: To address whether SP fish do not reproduce, we set up ten adult couples of SP males with LP females, as well as SP females with LP males. *Experiment 3*: To investigate the effect of photoperiod exposure on gonadotrope cell activity, fish were raised for 4 months in LP and exposed to SP, while those raised in SP for 4 months were exposed to LP, and sampled at day 0 (before exposure), 7, 14, and 30. As controls, the fish raised either in LP or in SP stays in their original photoperiod condition during the trial period.

### Euthanasia, morphometric measurements, and gonadosomatic index (GSI) calculation

The fish were euthanized by ice water immersion before the total body weight (mg) and the standard length (cm) were measured. Gonad weight (mg) was measured to calculate gonadosomatic index (GSI) using the following formula:

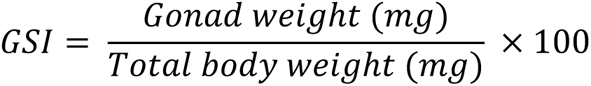

### Labelling of mitotic cells

To investigate the photoperiodic effect on gonadotrope mitosis, dTg fish that underwent *Experiment 3* were treated at day five with 1mM BrdU (Sigma) diluted in water with 0.33% DMSO for 4 h. Immunofluorescence was performed as previously described in (77) with minor modification. After an overnight fixation at 4 °C in 4% paraformaldehyde (PFA; Electron Microscopy 135 Sciences) in phosphate buffered saline with Tween (PBST: PBS, 0.1%; Tween-20), the brain and pituitary complex from dTg fish were washed three times with PBST, embedded in 3% agarose (H_2_O), and para-sagittally sectioned with 80 µm thickness using a vibratome (Leica). The free-floating sections were treated with 2M HCl at 37 °C for 1 hour for epitope retrieval before a 1-hour incubation at RT in blocking solution (3% normal goat serum (NGS); 0.5% Triton; 1% dimethylsulfoxide (DMSO) in PBST). The tissue slices were then incubated at 4 °C overnight with previously verified anti-BrdU antibody (abcam; ab6326) and anti-Fshβ antibody. After extensive washes with PBST, the slices were incubated for 4 hours at RT with secondary antibodies (**Supp. Table 1**).

### Melatonin and Tsh effects on dispersed pituitary cell culture (*in vitro*) and pituitary organ culture (*ex vivo*)

To investigate the direct effect of melatonin or Tsh, an *in vitro* dispersed pituitary cell culture experiment was performed as previously described (27). Cells from four pituitaries from adult WT fish were dissociated as previously described (78). Pituitary cells were mechanically dissociated using a glass pipette after a 30-minute digestion in trypsin (2 mg/ml; Sigma), followed by a 20-minute incubation in trypsin inhibitor (1 mg/ml; Sigma) and Dnase I type IV (2 µg/ml; Sigma). After centrifugation at 200 *g*, the cells were resuspended in 200 µl growth medium (L-15; Life Technologies) adjusted to 280-290 mOsm with mannitol and pH 7.75 with 1.8 mM glucose, 10 mM NaHCO_3_, and penicillin/streptomycin (50 U/ml; Lonza). The cells were plated in a 48-well plastic plate (Sarstedt) coated with poly-L-lysine (Sigma). The plate was prepared by adding 50 µl poly-L-lysine, left for 1 minute before decanting the solution and washed with 500 µl MQ water, and air drying the coated wells in a UV-light laminar flow hood for approximately 30 minutes. The experiment was run in triplicate.

To investigate the indirect effect of melatonin or Tsh, an *ex vivo* isolated pituitary organ culture was performed by detaching pituitaries from the brain and culturing three pituitaries together in growth medium supplemented with 5% fetal bovine serum (Sigma) as previously described (79). The medium for all experiments above was supplemented with 10 µM of melatonin (Sigma) or 0.5 µM bovine TSH pituitary extract (Sigma) or vehicle alone and incubated for 24 h at 26°C and 1% CO_2_. The TSH effect on gonadotrope cell mitosis was also evaluated using pituitary organ culture treated with 12 hours of TSH and 3 hours of BrdU, as explained above, before immunofluorescence (see Fig. 3C for illustration).

### Sex steroid extraction and enzyme-linked immunosorbent assays (ELISA)

Blood was sampled as previously described (80). Blood was collected from the caudal vein using a glass needle (outer diameter 1 mm, inner diameter 0.5 mm; GD-1; Narishige) coated with 0.05 U/µl heparin sodium (Sigma) in phosphate buffered saline (PBS). The blood was stored at –80 °C until use. Sex steroid extraction and ELISA were performed as previously described (81). Briefly, the blood samples were diluted 200× in PBS, extracted with diethyl ether (Sigma), and the E2 and 11-KT levels were measured using E2 and 11-KT ELISA kits (Cayman Chemical) according to the manufacturer’s instructions. Sex steroid concentrations were calculated with a standard curve fitted by a 4-parameter-logistic regression (R^2^ > 0.99).

### Bulk and single cell (sc) pituitary transcriptomics analysis (RNA-seq)

We used an available RNA-seq dataset from 16 male and 68 female medaka (82). Read counts per gene were normalized using the total amount of reads (counts per million).

For the scRNA-seq, we used an available RNA-seq dataset from 16 male and 68 female medaka (83). We used the Seurat (version 3.2.3, R version 4.0.3) function FindMarkers to determine which genes are specific for the cell cluster expressing tshr and aromatase. Using the default Wilcoxon rank sum test for genes expressed in at least two cells in that cluster relative to all other cells, and showing a minimum log fold change of 1, we found 90 genes with an adjusted *p*-value (Bonferroni corrected) of <0.05. Several of these genes are known as FS markers form literature on other teleost species.

### Quantification of mRNA levels with quantitative polymerase chain reaction (qPCR)

qPCR was performed as previously described in (84). For *in vivo* and *ex vivo* experiments, the pituitary was collected and stored at –80 °C in 300 µl of TRIzol (Invitrogen) with 6-7 zirconium oxide beads (Bertin Technologies). For *in vitro* experiments, 300 µl of TRIzol was added to the wells, which were scraped using pipette tips before transfer to the tubes and storage at –80 °C until use. The samples were homogenized and mixed with 120 µl chloroform, and the pellet was reconstituted with 14 µl nuclease free water. cDNA synthesis was performed with a minimum of 120 ng of RNA using SuperScript III Reverse Transcriptase (Invitrogen) and random hexamer primers (Thermofisher Scientific). cDNA samples (5× diluted) were analyzed in duplicate, using 3 µl of the cDNA and 5 µM each of forward and reverse primer in a total volume of 10 µl (**Supp. Table 2**). The parameter cycle was 10 min pre-incubation at 95 °C, followed by 42 cycles of 95 °C for 10 s, 60 °C for 10 s and 72 °C for 6 s, followed by melting curve analysis. The mRNA level was normalized using *gapdh* and *rpl7* as reference genes.

### Identification of the cells expressing Tsh receptors with RNAscope

RNAscope was performed as previously described (85). After euthanasia, the blood was removed by cardiac perfusion as described previously (86) with 4% PFA (PBS). Brain and pituitary were dissected and fixed overnight at 4 °C with 4% PFA (PBST). Tissues were then incubated in 25% sucrose solution (diluted in PBS) overnight at 4 °C and mounted in a block with OCT (Tissue-Tek, Sakura) and stored at –80 °C until use. Tissues were later parasagittally sectioned with a CM3050 Leica cryostat (10 µm sections). RNAscope fluorescent multiplex V2 assay (87) (ACDbio) was carried out as described by the supplier using *tshr2* (ENSORLG00000014222) and *cyp19a1b* (ENSORLG00000005548) probes and combined with opals 520 (*tshr2*) and 690 (*cyp19a1b*) (Akoya Bioscience) to avoid signal crosstalk during imaging.

### Labeling of folliculostellate cells

Pituitaries from adult dTg fish were directly incubated at RT in the dark for 4 hours in 100 µM dipeptide β-Ala-Lys-Nε-AMCA (US Biological Life Sciences) (88) diluted in Ca^2+^-free extracellular solution (2.9 mM KCl, 134 mM NaCl, 1.2 mM MgCl2, 4.5 mM glucose, and 10 mM N-2-hydroxyethylpiperazine-N-2-ethane sulfonic acid (HEPES), adjusted to osmolality 280-290 mOsm with mannitol and pH 7.75 with 1 M NaOH) with 0.5% bovine serum albumin. The pituitaries were then fixed in 4% PFA overnight and washed in PBST three times for 10 minutes at RT and incubated two days in PBST at 4 °C before mounting.

### Image processing and cell counting

Fluorescent images were obtained using a Leica Confocal Microscope (Dmi8, Leica) with 20× apochromat objective (numerical aperture 0.75), with laser wavelength 488 (Alexa-488, Opal 520), 555 (Alexa-555), 647 (Alexa-647, Opal 690). To avoid cross talk between fluorophores, the channels were obtained sequentially. Las X (v3.7, Leica) and ImageJ (1.53t; http://rsbweb.nih.gov/ij/) were used to process the images. Automatic cell counting was performed according to (28) while triple-labelled cells were counted manually using an ImageJ-based cell-counter plugin.

### Statistical analysis

Data were tested for normality and homogeneity with Shapiro-Wilk and Levene’s tests, respectively. Differences in mRNA levels and cell number were evaluated by independent-sample Student’s t-test or Mann-Whitney U test for non-parametric analysis. Statistical significance was set to *p* < 0.05. All statistical analyses were performed using Jamovi (Version 2.2.5) (89), and the graphs are provided as mean ± standard error of the mean (SEM).

## DECLARATIONS

### Ethics approval

Animal experiments were performed according to the regulation of the care and welfare of research animals in Norway and at the Norwegian University of Life Sciences.

### Data availability

All datasets generated during and/or analyzed during the current study are present in this manuscript or have been previously published.

## FUNDING

This work was funded by the Norwegian University of Life Sciences (to RF), by the Research Council of Norway (grant numbers 244461 and 243811, Aquaculture program to FAW), and Fiskeri – og havbruksnæringens forskningsfond (grant number 901590 to KH).

## ACKNOWLEDGEMENTS

We thank Anthony Peltier, Lourdes Carreon G Tan, and Arturas Kavaliauskis for fish facility maintenance. We also than Prof. Dianne Baker for the English language editing.

## FIGURES AND TABLES

**Supp. Fig. 1.**
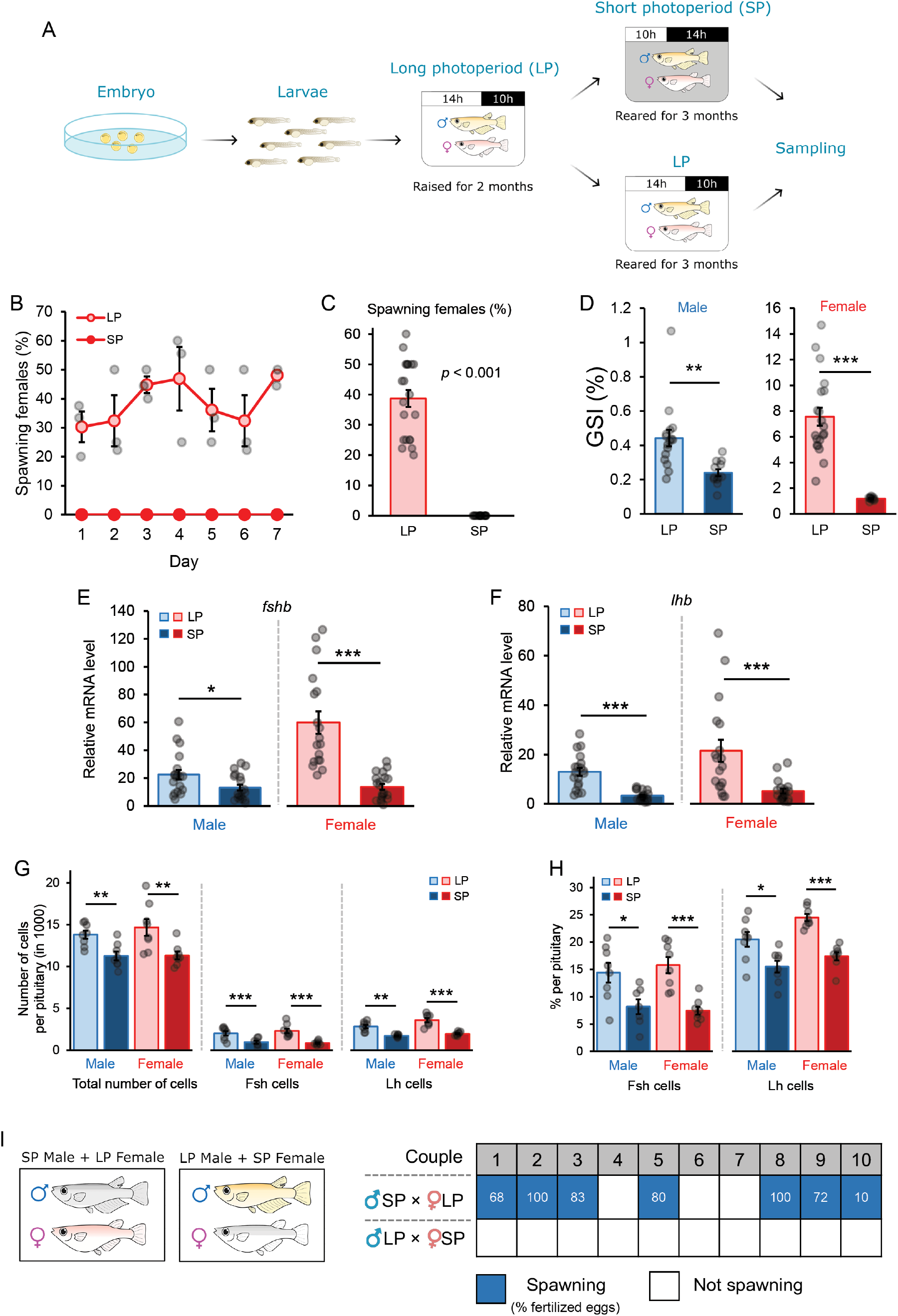
Long photoperiod stimulates reproduction by increasing gonadotrope cell activity (hormone synthesis) and cell number. (A) Illustration of the experimental protocol used. Two-month-old medaka fish raised in long photoperiod (LP) were divided into two groups reared under long (LP) and short (SP) photoperiod for 3 months before sampling. (B) Graph showing the daily percentage of spawning females kept in SP and LP during 7-day observation (n = 3). (C) Graph showing the average daily percentage of spawning females kept in SP and LP for 7 consecutive days (n = 21). (D) Gonadosomatic index (GSI) in male and female medaka kept in SP and LP (n = 16-20). (E-F) Graph presenting the relative mRNA levels of *fshb* and *lhb* in the pituitary of male and female medaka in SP and LP (n = 19-20). (G-H) Graph presenting the absolute (G) and relative (H) number of gonadotrope cells per pituitary in SP and LP fish (n = 7-8). (I) Spawning observation in SP fish coupled with LP fish to evaluate reproductive capacity of SP fish. Number in the square indicates the percentage of fertilized eggs. Statistical analyses were performed using two-sample independent t-test or Mann Whitney U test, in which the graph was provided as mean ± SEM while the jittered dots represent each individual (* < 0.05; ** < 0.01; *** < 0.001).

**Supp. Fig. 2:**
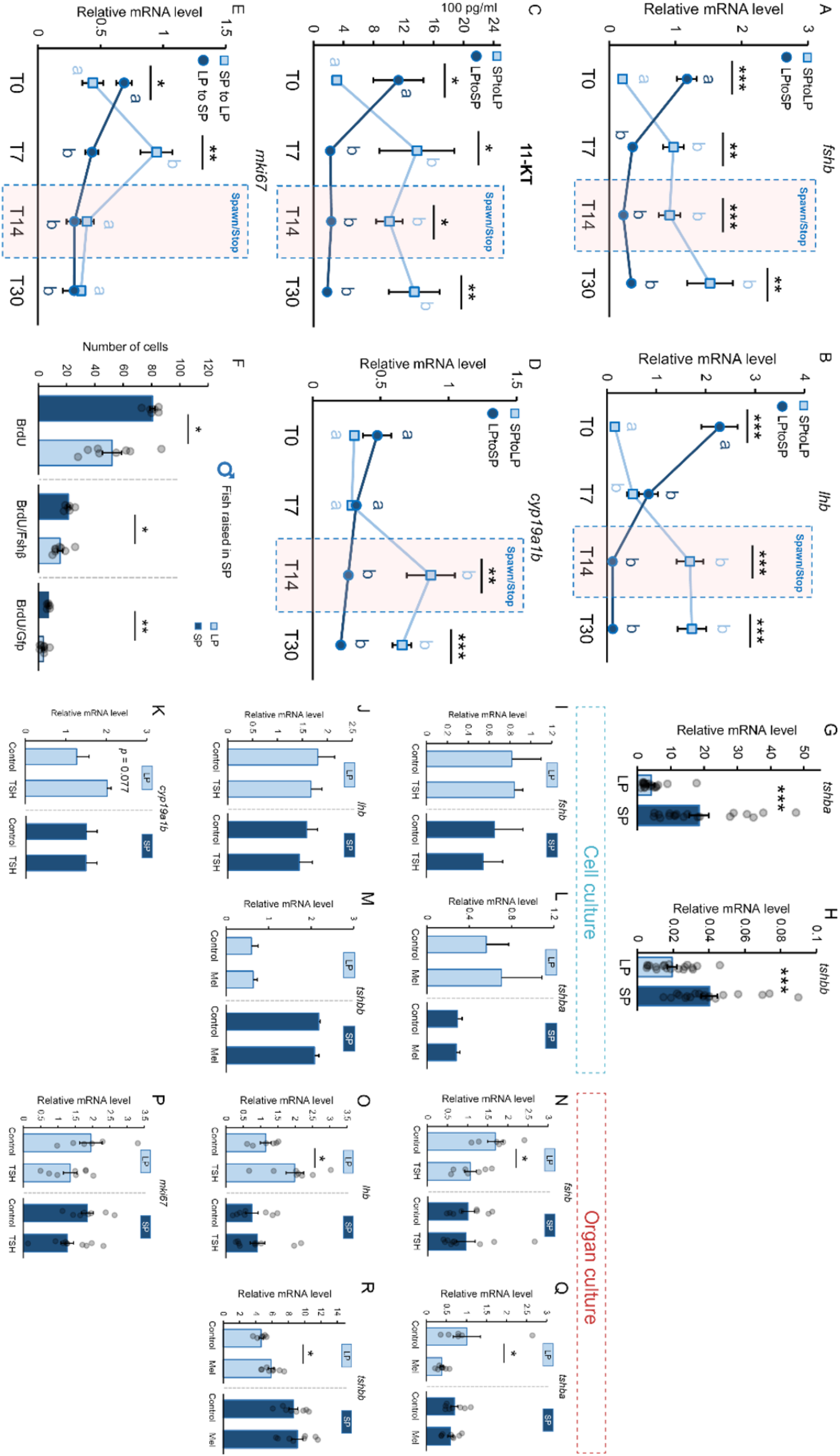
Efffect of photoperiod, Tsh and melatonin in males. (A-E) The fluctuation of *fshb*, *lhb*, *cyp19a1b*, and *mki67* expression as well as 11-KT levels observed in males at different time points following the photoperiod change (n = 6-11). Black asterisks display statistically differences between the SPtoLP group and the SPtoLP group for each observation day. The letters (a, b) show statistically differences between each time points and T0 which is taken as reference within one treatment group. The statistical analyses were performed using two-sample independent t-test or Mann-Whitney U test (* < 0.05; ** < 0.01; *** < 0.001). Like in females, *fshb* and *lhb* increase in SPtoLP while decrease in LPtoSP males. Significant changes are already observed for both *fshb* and *lhb* after 7 days. In line with gonadotropin mNRA levels, 11-keto testosterone (11-KT) levels also significantly change in 7 days, increasing in SPtoLP while decreasing in LPtoSP males. Also similar to in females, the mRNA levels for aromatase (*cyp19a1b*) show the same pattern that gonadotropins and sex steroids levels, suggesting that the same mechanisms take place in males and females. While *mki67* levels increase in SPtoLP males, the total number of mitotic cells as well as the number of both Lh and Fsh gonadoaropes dividing (F) in SP fish exposed to LP for 5 days (n = 6-8) cell, are lower in males moved from SP to LP than in males kept in SP. These results suggest that opposite to in females, LP inhibits gonadotrope cell proliferation in males. Because of the number of gonadotropes and percentage of cells they represent in the pituitary is higher in LP males than in SP males, our results therefore suggest than another mechanism is playing a role in gonadotrope cell proliferation in males under LP conditions. *tshba* (G) and *tshbb* (H) mRNA levels in LP and SP condition in male medaka (n = 19-20). As in females, *tshbb* levels are reduced in LP males compared to SP males. In contrast, the levels of *tshba* show sexual dimorphism with LP males showing lower *tshba* levels than SP males. Effect of 0.5 µM bovine TSH pituitary extract, 10 µM melatonin, or vehicle (control) on *fshb*, *lhb*, *cyp19a1b*, *tshba*, and *tshbb* mRNA levels in dispersed pituitary cell cultures (I-M) or *ex vivo* medaka pituitary organ cultures (N-R) in male medaka from LP and SP condition (n = 3; in which each replicate represents 4 pooled pituitaries for *in vitro* studies and n = 6-10 for *ex vivo* studies). The statistical analyses were performed using two-sample independent t-test or Mann Whitney U test. All graphs (unless otherwise stated) are represented as mean ± SEM with the jitter dots representing each individual (* < 0.05; *** < 0.001). We found that similar to in females, TSH has an effects on gonadotropin mRNA levels in *ex vivo* pituitary cultures but not in dissociated pituitary cell cultures. Surprisingly, TSH stimulates *lhb* but inhibits *fshb* levels. Interestingly, the effects on gonadotropins can only be observed in LP males, and not SP males. Aromatase mRNA levels increased at nearly significance levels in LP male cell cultures following TSH stimulation. Together, these results suggest that, as in females, TSH regulate gonadotropin mNRA synthesis in males, but in an opposite manner than females for *fshb*. As in females, while no effects were observed in dissociated cell cultures, *tshba* and *tshbb* levels were regulated by melatonin in *ex vivo* pituitary organ cultures. Also similar to in females, melatonin increased *tshbb* and reduced *tshba* levels in LP males. It is surprising that, melatonin inhibits *tshba* in males, as SP males which are expected to have higher melatonin levels were found to possess higher *tshba* levels than LP males. These results thus suggest that another factor plays a more important role in the regulation of *tshba* in SP males. Nevertheless, the higher levels of *tshba* in SP males (compared to in LP males), are in line with the inhibitory effect of TSH observed on *fshb* and the lower levels of *fshb* observed in SP males (compared to in LP males). Finally, in contrast to in females, TSH had no effects on *mki67* levels, supporting the hypothesis that photoperiod does not regulate gonadotrope cell proliferation via Tsh in the male pituitary, and thus that another mechanism is taking place.

**Supp. Fig. 3.**
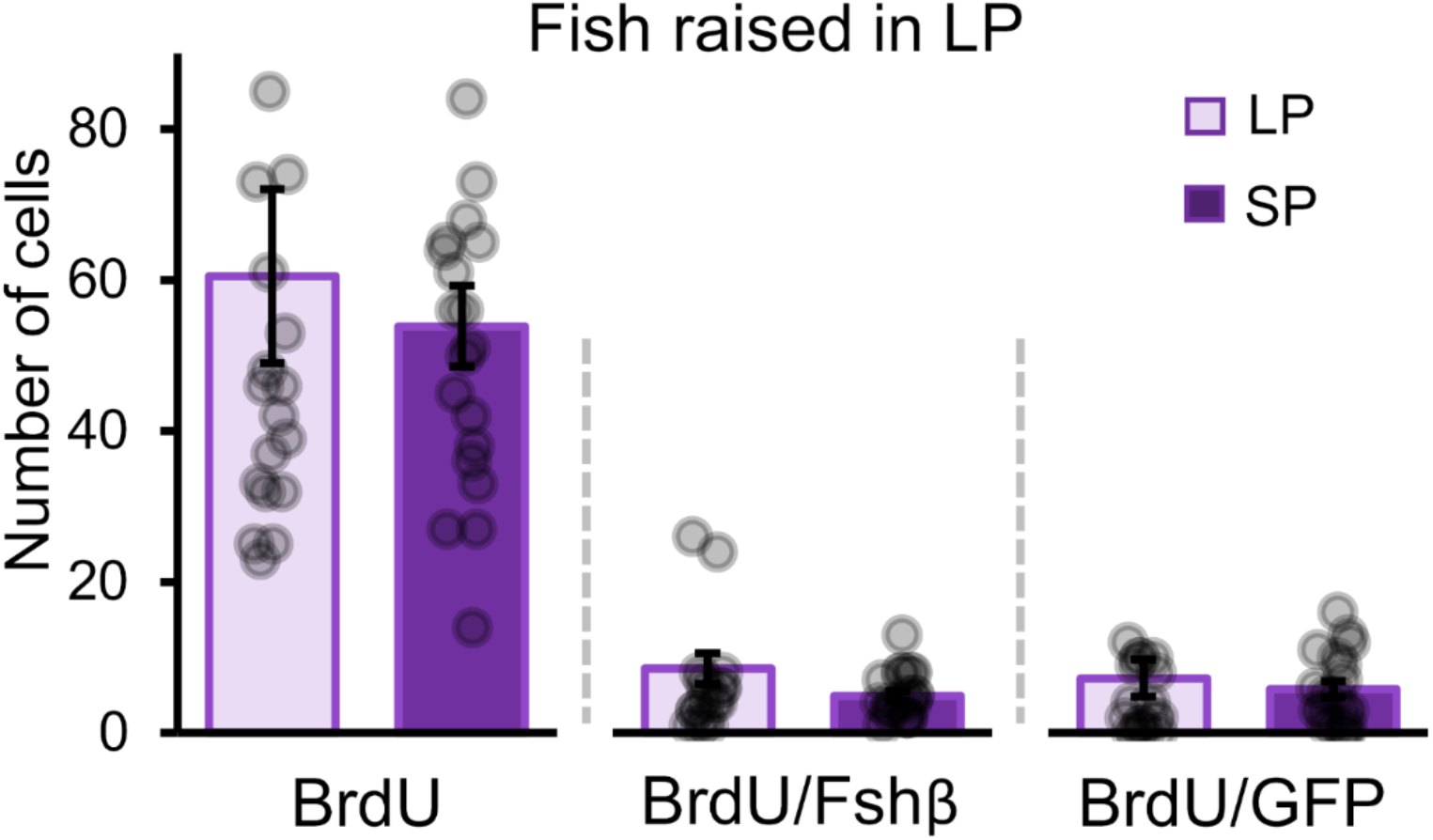
No change in the number of mitotic cells when LP raised fish exposed to SP. Number of mitotic cells 5 days after the photoperiod regime change in mixed-sex groups (n = 19-20). The statistical analyses were performed using two-sample independent t-test, in which the graph represents mean ± SEM while the jittered dots represent each individual.

**Supp. Fig. 4.**
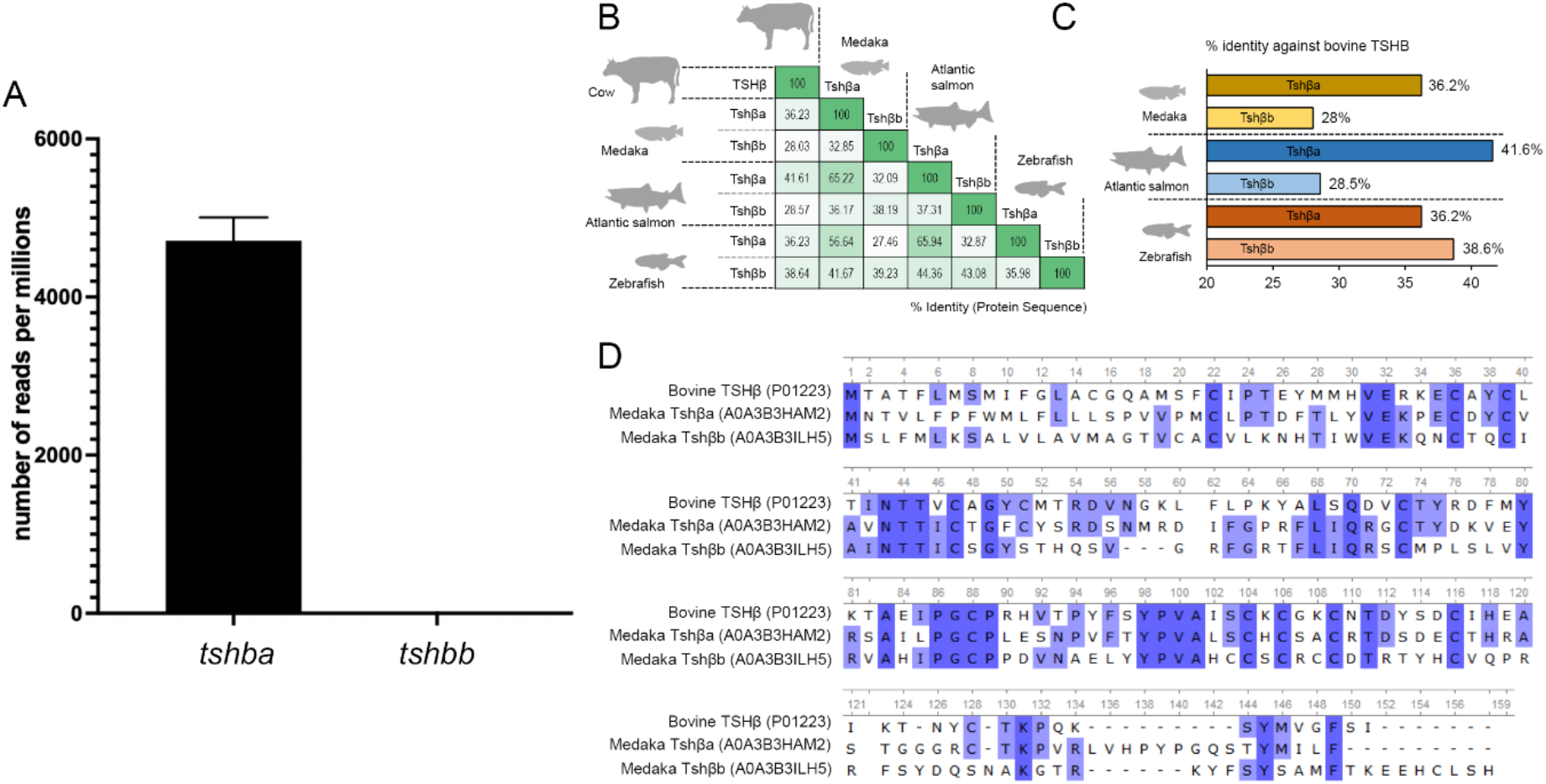
Protein sequence identity of bovine TSHβ shows a more similarity to medaka Tshβa than Tshβb. (A) Number of reads of *tshba* and *tshbb* in from the medaka pituitary RNAseq dataset from both sexes combined. (B) Percentage of protein sequence identity of bovine TSHβ (P01223) and other fish species, including medaka (*Oryzias latipes*; Tshβa: A0A3B3HAM2; Tshβb: A0A3B3ILH5), Atlantic salmon (*Salmo salar*; Tshβa: O73824; Tshβb: A0A1S3MIK7), and zebrafish (*Danio rerio*, Tshβa: A0A8M9PLW7; Tshβb: A0A8M6Z4G8). (C) Percentage of protein sequence identity of bovine TSHβ to medaka, Atlantic salmon, and zebrafish Tshβa and Tshβb. (D) Sequence alignment of medaka Tshβa and Tshβb to bovine TSHβ as the reference sequence showing that even if some regions seem well conserved between the mammalian TSHβ and the medaka Tshβa and Tshβb protein sequences, they still are strongly divergent. Sequences were retrieved from UniProt, while percentage of sequence identity and alignment was performed using default settings in UniProt using Clustal Omega algorithm.

**Supp. Fig. 5.**
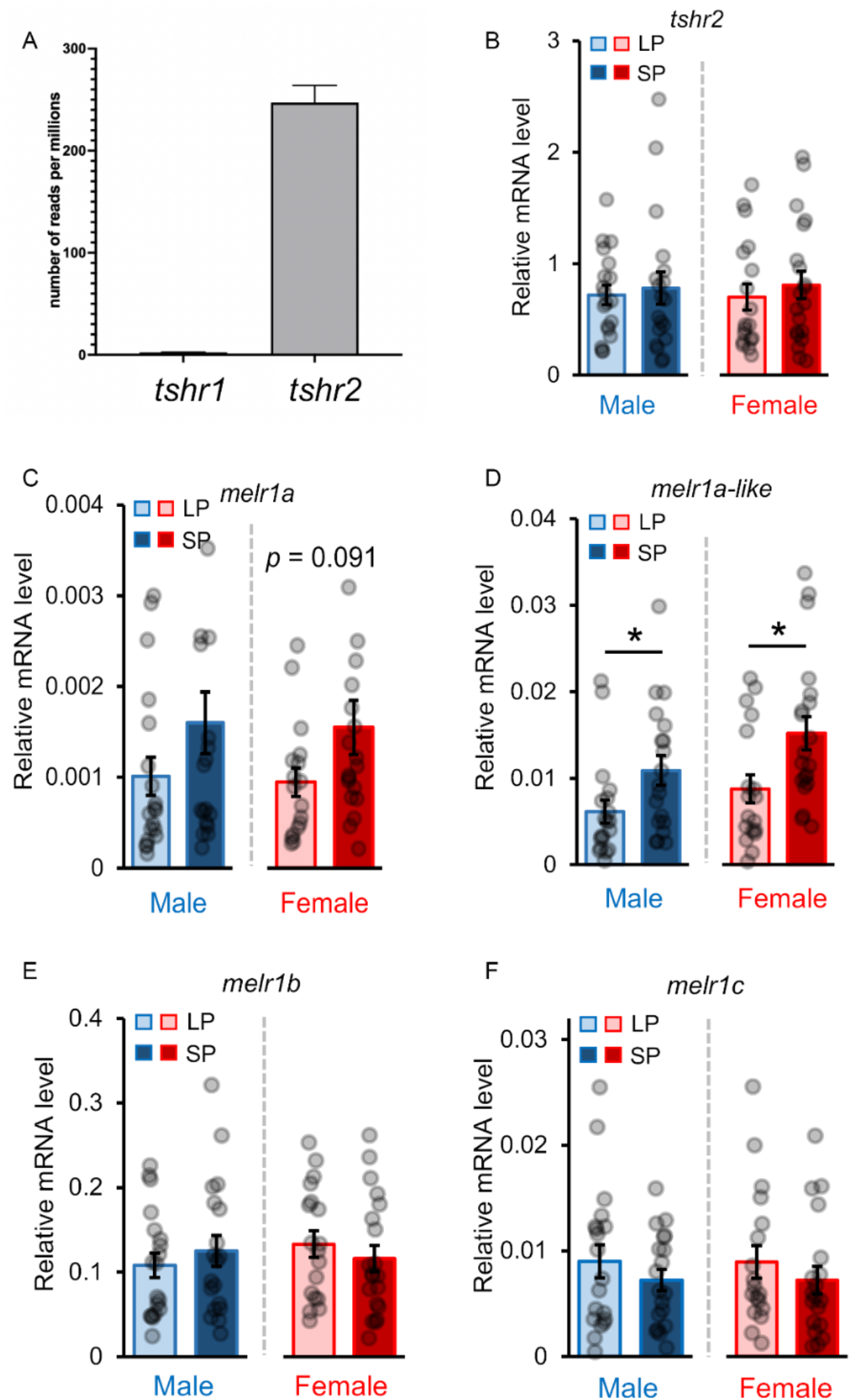
One of the two identified Tsh receptors (*tshr2*) is expressed in the medaka pituitary with photoperiod-independent expression. (A) Number of reads per million of two Tsh receptors (arbitrarily named *tshr1* and *tshr2)* in the medaka pituitary from RNA transcriptomic data from both sexes combined. (B) The mRNA levels of *tshr2* in the male and female medaka pituitary in SP and LP condition (n = 19-20). (C-D) The relative mRNA levels four melatonin receptor paralogs in the medaka pituitary, *melr1a* (A), *melr1a-like* (B), *melr1b* (C), and *melr1c* (D) from SP and LP condition (n = 19-20). The statistical analyses were performed using two-sample independent t-test or Mann Whitney U test, in which the graph represents mean ± SEM while jitter dots represent each individual (* < 0.05).

**Supp. Fig. 6.**
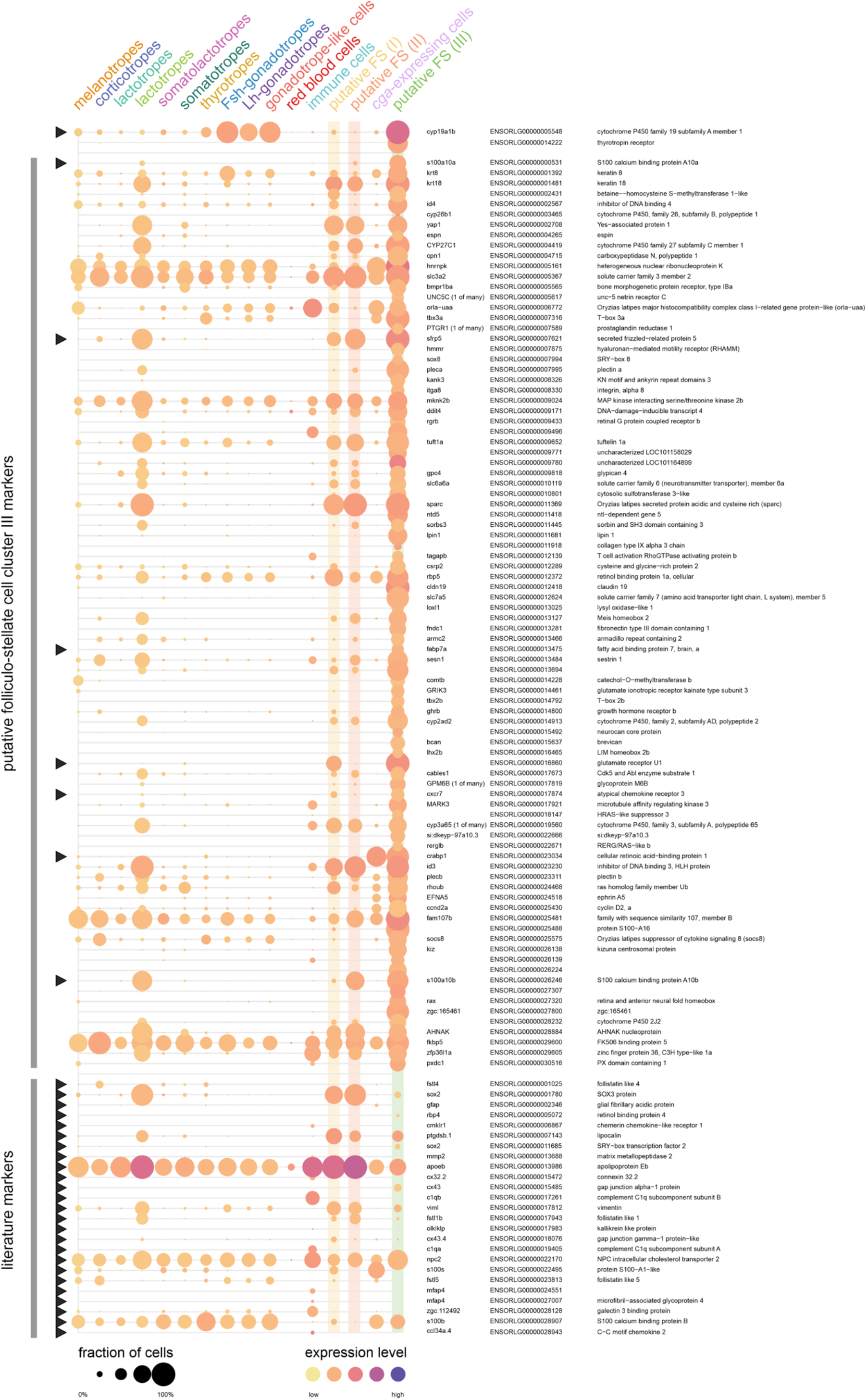
Genes specifically expressed in putative folliculo-stellate cell cluster III. Expression levels are indicated by colour, the percentage of number of cells in a cluster expressing a gene by circle size. At the top, the expression patterns of tshr2 and cyp19a1b summarize Figure 5 B-C. Below that, an additionall 88 genes are specifically expressed in the same cells. This set includes several previously found marker genes of either folliculostellate cells* or pituicytes# (black triangles). At the bottom, homologues of other marker genes do not provide a clear identification of folliculo-stellate cells in the medaka pituitary.

**Supp. Table 1.** Primary and secondary antibodies used for immunofluorescence.

| Antibody | Dilution | Source | Reference |
| --- | --- | --- | --- |
| Rat anti-BrdU | 1 : 250 | abcam (ab6326) | (28) |
| Rabbit anti-medakaFshβ | 1 : 500 | custom-made | (84) |
| Goat anti-rabbit (Alexa-555) | 1 : 500 | Invitrogen (A21429) | (35) |
| Donkey anti-rat (Alexa-647) | 1 : 500 | Invitrogen (A78947) | (28) |

**Supp. Table 2.**
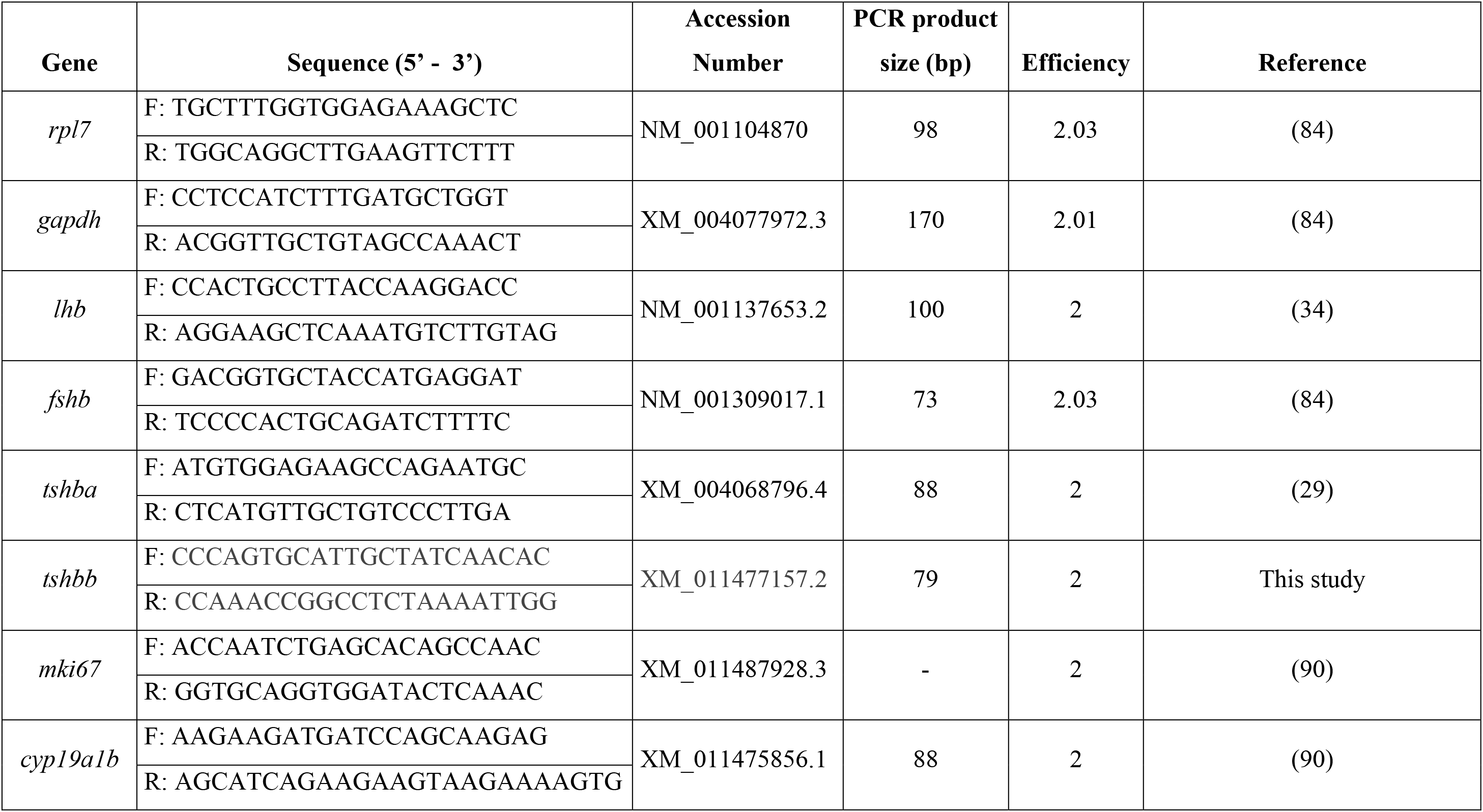
Primer sequences used for mRNA level analysis in the medaka pituitary.

## Notes

### Competing Interest Statement

The authors have declared no competing interest.

## REFERENCES

1. A. Shinomiya, T. Shimmura, T. Nishiwaki-Ohkawa, T. Yoshimura, Regulation of Seasonal Reproduction by Hypothalamic Activation of Thyroid Hormone. Frontiers in Endocrinology 5 (2014).

2. F. H. Bronson, Climate change and seasonal reproduction in mammals. Philos Trans R Soc Lond B Biol Sci 364, 3331–3340 (2009).

3. H. Migaud, A. Davie, J. F. Taylor, Current knowledge on the photoneuroendocrine regulation of reproduction in temperate fish species. Journal of Fish Biology 76, 27–68 (2010).

4. S. Kanda, Evolution of the regulatory mechanisms for the hypothalamic-pituitary-gonadal axis in vertebrates–hypothesis from a comparative view. General and Comparative Endocrinology 284, 113075 (2019).

5. Z. Yaron, B. Levavi-Sivan, “Hormonal control of reproduction and growth | Endocrine Regulation of Fish Reproduction” in Encyclopedia of Fish Physiology, A. P. Farrell, Ed. (Academic Press, San Diego, 2011), vol. 2, pp. 1500–1508.

6. R. Fontaine, M. R. Royan, K. von Krogh, F.-A. Weltzien, D. M. Baker, Direct and Indirect Effects of Sex Steroids on Gonadotrope Cell Plasticity in the Teleost Fish Pituitary. Frontiers in Endocrinology 11 (2020).

7. D. Zambrano, Innervation of the teleost pituitary. General and Comparative Endocrinology 3, 22–31 (1972).

8. F.-A. Weltzien, J. Hildahl, K. Hodne, K. Okubo, T. M. Haug, Embryonic development of gonadotrope cells and gonadotropic hormones – Lessons from model fish. Molecular and Cellular Endocrinology 385, 18–27 (2014).

9. R. Fontaine et al., Pituitary multi-hormone cells in mammals and fish: history, origin, and roles. Frontiers in Neuroendocrinology 67, 101018 (2022).

10. E. Tolla, T. J. Stevenson, Sex Differences and the Neuroendocrine Regulation of Seasonal Reproduction by Supplementary Environmental Cues. Integrative and Comparative Biology 60, 1506–1516 (2020).

11. E. Ciani et al., Effects of Melatonin on Anterior Pituitary Plasticity: A Comparison Between Mammals and Teleosts. Frontiers in Endocrinology 11 (2021).

12. Y. Nakane, T. Yoshimura, Universality and diversity in the signal transduction pathway that regulates seasonal reproduction in vertebrates. Frontiers in Neuroscience 8 (2014).

13. Y. Nakane, T. Yoshimura, Photoperiodic Regulation of Reproduction in Vertebrates. Annual Review of Animal Biosciences 7, 173–194 (2019).

14. H. Dardente, V. Simonneaux, GnRH and the photoperiodic control of seasonal reproduction: Delegating the task to kisspeptin and RFRP-3. Journal of Neuroendocrinology 34, e13124 (2022).

15. V. Simonneaux, A Kiss to drive rhythms in reproduction. European Journal of Neuroscience 51, 509–530 (2020).

16. A. Acharjee, R. Chaube, K. P. Joy, Effects of altered photoperiod and temperature on expression levels of gonadotrophin subunit mRNAs in the female stinging catfish Heteropneustes fossilis. Journal of Fish Biology 90, 2289–2311 (2017).

17. C. H. Lee, Y. J. Park, Y. D. Lee, Effects of Photoperiod Manipulation on Gonadal Activity of the Damselfish, Chromis notata. Dev Reprod 21, 223–228 (2017).

18. K. Okuzawa, K. Furukawa, K. Aida, I. Hanyu, Effects of photoperiod and temperature on gonadal maturation, and plasma steroid and gonadotropin levels in a cyprinid fish, the honmoroko Gnathopogon caerulescens. Gen Comp Endocrinol 75, 139–147 (1989).

19. K. Tsuneki, A Systematic Survey of the Occurrence of the Hypothalamic Saccus Vasculosus in Teleost Fish. Acta Zoologica 73, 67–77 (1992).

20. Y. Nakane et al., The saccus vasculosus of fish is a sensor of seasonal changes in day length. Nature Communications 4, 2108 (2013).

21. M. S. Fleming et al., Functional divergence of thyrotropin beta-subunit paralogs gives new insights into salmon smoltification metamorphosis. Scientific Reports 9, 4561 (2019).

22. S. Irachi et al., Photoperiodic regulation of pituitary thyroid-stimulating hormone and brain deiodinase in Atlantic salmon. Molecular and Cellular Endocrinology 519, 111056 (2021).

23. G. Maugars, S. Dufour, J. Cohen-Tannoudji, B. Quérat, Multiple Thyrotropin β-Subunit and Thyrotropin Receptor-Related Genes Arose during Vertebrate Evolution. PLOS ONE 9, e111361 (2014).

24. K. Sakai, Y. Yamamoto, T. Ikeuchi, Vertebrates originally possess four functional subtypes of G protein-coupled melatonin receptor. Scientific Reports 9, 9465 (2019).

25. G. Maugars, R. Nourizadeh-Lillabadi, F.-A. Weltzien, New Insights Into the Evolutionary History of Melatonin Receptors in Vertebrates, With Particular Focus on Teleosts. Frontiers in Endocrinology 11 (2020).

26. E. Ciani et al., Melatonin receptors in Atlantic salmon stimulate cAMP levels in heterologous cell lines and show season-dependent daily variations in pituitary expression levels. Journal of Pineal Research 67, e12590 (2019).

27. R. Fontaine, E. Ager-Wick, K. Hodne, F.-A. Weltzien, Plasticity in medaka gonadotropes via cell proliferation and phenotypic conversion. Journal of Endocrinology 245, 21 (2020).

28. R. Fontaine, E. Ager-Wick, K. Hodne, F.-A. Weltzien, Plasticity of Lh cells caused by cell proliferation and recruitment of existing cells. Journal of Endocrinology 240, 361 (2019).

29. M. R. Royan et al., 3D Atlas of the Pituitary Gland of the Model Fish Medaka (Oryzias latipes). Frontiers in Endocrinology 12 (2021).

30. R. Fontaine et al., Gonadotrope plasticity at cellular, population and structural levels: A comparison between fishes and mammals. General and Comparative Endocrinology 287, 113344 (2020).

31. J. F. F. Powell, S. L. Krueckl, P. M. Collins, N. M. Sherwood, Molecular forms of GnRH in three model fishes: rockfish, medaka and zebrafish. 150, 17 (1996).

32. J. Wittbrodt, A. Shima, M. Schartl, Medaka — a model organism from the far east. Nature Reviews Genetics 3, 53–64 (2002).

33. H. Hori, “A Glance at the Past of Medaka Fish Biology” in Medaka: A Model for Organogenesis, Human Disease, and Evolution, K. Naruse, M. Tanaka, H. Takeda, Eds. (Springer Japan, Tokyo, 2011), https://doi.org/10.1007/978-4-431-92691-7_1, pp. 1–16.

34. J. Hildahl et al., Developmental tracing of luteinizing hormone β-subunit gene expression using green fluorescent protein transgenic medaka (Oryzias latipes) reveals a putative novel developmental function. 241, 1665–1677 (2012).

35. K. Hodne, R. Fontaine, E. Ager-Wick, F.-A. Weltzien, Gnrh1-Induced Responses Are Indirect in Female Medaka Fsh Cells, Generated Through Cellular Networks. Endocrinology 160, 3018–3032 (2019).

36. E. Ager-Wick et al., Integrative transcriptomics reveals ectopic lipid homeostasis mechanisms in non-endocrine cells of the teleost pituitary. bioRxiv 10.1101/2021.06.11.448009, 2021.2006.2011.448009 (2022).

37. X. d. A. d. Tassigny, W. H. Colledge, The Role of Kisspeptin Signaling in Reproduction. Physiology 25, 207–217 (2010).

38. Y. Kawabata-Sakata, Y. Nishiike, T. Fleming, Y. Kikuchi, K. Okubo, Androgen-dependent sexual dimorphism in pituitary tryptophan hydroxylase expression: relevance to sex differences in pituitary hormones. Proceedings of the Royal Society B: Biological Sciences 287, 20200713 (2020).

39. C. S. Koger, S. J. Teh, D. E. Hinton, Variations of Light and Temperature Regimes and Resulting Effects on Reproductive Parameters in Medaka (Oryzias latipes)1. Biology of Reproduction 61, 1287–1293 (1999).

40. D. N. Weber, R. E. Spieler, Effects of the light-dark cycle and scheduled feeding on behavioral and reproductive rhythms of the cyprinodont fish, Medaka,Oryzias latipes. Experientia 43, 621–624 (1987).

41. A. Hellqvist, M. Schmitz, B. Borg, Effects of castration and androgen-treatment on the expression of FSH-β and LH-β in the three-spine stickleback, gasterosteus aculeatus—Feedback differences mediating the photoperiodic maturation response? General and Comparative Endocrinology 158, 178–182 (2008).

42. Y. T. Shao et al., Androgen feedback effects on LH and FSH, and photoperiodic control of reproduction in male three-spined sticklebacks, Gasterosteus aculeatus. General and Comparative Endocrinology 182, 16–23 (2013).

43. W. Li et al., Effects of different photoperiod conditions on survival, growth, and gonadal development of Takifugu rubripes adults. Aquaculture 564, 739048 (2023).

44. E. L. Bittman, A. E. Jetton, C. Villalba, G. J. Devries, Effects of photoperiod and androgen on pituitary function and neuropeptide staining in Siberian hamsters. *American Journal of Physiology-Regulatory*, Integrative and Comparative Physiology 271, R64–R72 (1996).

45. M. R. Royan, D. Kayo, F.-A. Weltzien, R. Fontaine, Sexually dimorphic regulation of gonadotrope cell hyperplasia in medaka pituitary via mitosis and transdifferentiation. Endocrinology https://doi.org/10.1210/endocr/bqad030 (2023).

46. A. Takahashi, S. Kanda, T. Abe, Y. Oka, Evolution of the Hypothalamic-Pituitary-Gonadal Axis Regulation in Vertebrates Revealed by Knockout Medaka. Endocrinology 157, 3994–4002 (2016).

47. N. Murozumi et al., Loss of Follicle-Stimulating Hormone Receptor Function Causes Masculinization and Suppression of Ovarian Development in Genetically Female Medaka. Endocrinology 155, 3136–3145 (2014).

48. T. Kitano et al., Roles of Gonadotropin Receptors in Sexual Development of Medaka. http://dx.doi.org/10.3390/cells11030387.

49. Z. Zhang, B. Zhu, W. Ge, Genetic Analysis of Zebrafish Gonadotropin (FSH and LH) Functions by TALEN-Mediated Gene Disruption. Molecular Endocrinology 29, 76–98 (2015).

50. W.-K. So, H.-F. Kwok, W. Ge, Zebrafish Gonadotropins and Their Receptors: II. Cloning and Characterization of Zebrafish Follicle-Stimulating Hormone and Luteinizing Hormone Subunits—Their Spatial-Temporal Expression Patterns and Receptor Specificity1. Biology of Reproduction 72, 1382–1396 (2005).

51. L. Chu, J. Li, Y. Liu, C. H. K. Cheng, Gonadotropin Signaling in Zebrafish Ovary and Testis Development: Insights From Gene Knockout Study. Molecular Endocrinology 29, 1743–1758 (2015).

52. J. N. Anasti, M. R. Flack, J. Froehlich, L. M. Nelson, B. C. Nisula, A potential novel mechanism for precocious puberty in juvenile hypothyroidism. The Journal of Clinical Endocrinology & Metabolism 80, 276–279 (1995).

53. G. Csaba, M. A. Shahin, O. Dobozy, Effect of combined gonadotropin-thyrotropin treatment on development of testis and ovarium in the chickling. Acta Physiol Acad Sci Hung 55, 163–168 (1980).

54. M. Migaud, M. Batailler, D. Pillon, I. Franceschini, B. Malpaux, Seasonal Changes in Cell Proliferation in the Adult Sheep Brain and Pars Tuberalis. Journal of Biological Rhythms 26, 486–496 (2011).

55. Shona H. Wood et al., Binary Switching of Calendar Cells in the Pituitary Defines the Phase of the Circannual Cycle in Mammals. Current Biology 25, 2651–2662 (2015).

56. T. Lucon-Xiccato et al., Medaka as a model for seasonal plasticity: Photoperiod-mediated changes in behaviour, cognition, and hormones. Hormones and Behavior 145, 105244 (2022).

57. M. Daimon, T. Katsumura, H. Sakamoto, S. Ansai, H. Takeuchi, Mating experiences with the same partner enhanced mating activities of naïve male medaka fish. Scientific Reports 12, 19665 (2022).

58. H. Ohga, K. Ohta, M. Matsuyama, Long-day stimulation increases thyroid-stimulating hormone expression and affects gonadal development in chub mackerel. Comparative Biochemistry and Physiology Part A: Molecular & Integrative Physiology 275, 111334 (2023).

59. S. Anbalagan et al., Pituicyte Cues Regulate the Development of Permeable Neuro-Vascular Interfaces. Dev Cell 47, 711–726 e715 (2018).

60. M. Golan, L. Hollander-Cohen, B. Levavi-Sivan, Stellate Cell Networks in the Teleost Pituitary. Sci Rep 6, 24426 (2016).

61. M. F. Prummel et al., Expression of the Thyroid-Stimulating Hormone Receptor in the Folliculo-Stellate Cells of the Human Anterior Pituitary. The Journal of Clinical Endocrinology & Metabolism 85, 4347–4353 (2000).

62. M. Theodoropoulou et al., Thyrotrophin receptor protein expression in normal and adenomatous human pituitary. Journal of Endocrinology 167, 7–13 (2000).

63. L. J. S. Brokken, O. Bakker, W. M. Wiersinga, M. F. Prummel, Functional Thyrotropin Receptor Expression in the Pituitary Folliculo-Stellate Cell Line TtT/GF. Exp Clin Endocrinol Diabetes 113, 13–20 (2005).

64. S. V. Grommen, S. Geysens, V. M. Darras, B. De Groef, Chicken folliculo-stellate cells express thyrotropin receptor mRNA. Domestic Animal Endocrinology 37, 236–242 (2009).

65. C. Denef, Paracrinicity: The Story of 30 Years of Cellular Pituitary Crosstalk. Journal of Neuroendocrinology 20, 1-70 (2008).

66. T. Fauquier, N. C. Guérineau, R. A. McKinney, K. Bauer, P. Mollard, Folliculostellate cell network: A route for long-distance communication in the anterior pituitary. Proceedings of the National Academy of Sciences 98, 8891 (2001).

67. W. Allaerts, H. Vankelecom, History and perspectives of pituitary folliculo-stellate cell research. European Journal of Endocrinology eur j endocrinol 153, 1–12 (2005).

68. P. R. Le Tissier, P. Mollard, Renewing an old interest: Pituitary folliculostellate cells. Journal of Neuroendocrinology 33, e13053 (2021).

69. T. Soji, D. C. Herbert, Intercellular communication between rat anterior pituitary cells. The Anatomical Record 224, 523–533 (1989).

70. D. J. Hodson et al., Existence of long-lasting experience-dependent plasticity in endocrine cell networks. Nature Communications 3, 605 (2012).

71. I. Morand et al., Cell-to-cell communication in the anterior pituitary: evidence for gap junction-mediated exchanges between endocrine cells and folliculostellate cells. Endocrinology 137, 3356–3367 (1996).

72. K. Sakai, Y. Yamamoto, T. Ikeuchi, Vertebrates originally possess four functional subtypes of G protein-coupled melatonin receptor. Sci Rep 9, 9465 (2019).

73. K. Sato, K. N. Nwe;, H. Ohuchi, The Opsin 3/Teleost multiple tissue opsin system: mRNA localization in the retina and brain of medaka (Oryzias latipes). Journal of Comparative Neurology 529, 2484–2516 (2021).

74. K. Sato et al., Two UV-Sensitive Photoreceptor Proteins, Opn5m and Opn5m2 in Ray-Finned Fish with Distinct Molecular Properties and Broad Distribution in the Retina and Brain. PLOS ONE 11, e0155339 (2016).

75. K. Fujisawa et al., Seasonal variations in photoperiod affect hepatic metabolism of medaka (Oryzias latipes). FEBS Open Bio 11, 1029–1040 (2021).

76. M. Kinoshita, K. Murata, K. Naruse, M. Tanaka, “Looking at Adult Medaka” in Medaka: Biology, Management, and Experimental Protocols, M. Kinoshita, K. Murata, K. Naruse, M. Tanaka, Eds. (Wiley Blackwell, 2009), https://doi.org/10.1002/9780813818849.ch5 chap. 5, pp. 117–164.

77. R. Fontaine et al., Dopamine Inhibits Reproduction in Female Zebrafish (Danio rerio) via Three Pituitary D2 Receptor Subtypes. Endocrinology 154, 807–818 (2013).

78. E. Ager-Wick et al., Preparation of a High-quality Primary Cell Culture from Fish Pituitaries. Journal of Visualized Experiments https://dx.doi.org/10.3791/58159, e58159 (2018).

79. T. Karigo et al., Time-of-Day-Dependent Changes in GnRH1 Neuronal Activities and Gonadotropin mRNA Expression in a Daily Spawning Fish, Medaka. Endocrinology 153, 3394–3404 (2012).

80. M. R. Royan et al., Gonadectomy and Blood Sampling Procedures in the Small Size Teleost Model Japanese Medaka (Oryzias latipes). JoVE https://dx.doi.org/10.3791/62006, e62006 (2020).

81. D. Kayo, Y. Oka, S. Kanda, Examination of methods for manipulating serum 17β-Estradiol (E2) levels by analysis of blood E2 concentration in medaka (Oryzias latipes). General and Comparative Endocrinology 285, 113272 (2020).

82. E. Ager-Wick et al., An RNA-seq time series of the medaka pituitary gland during sexual maturation. Scientific Data 10, 62 (2023).

83. K. Siddique, E. Ager-Wick, R. Fontaine, F.-A. Weltzien, C. V. Henkel, Characterization of hormone-producing cell types in the teleost pituitary gland using single-cell RNA-seq. Scientific Data 8, 279 (2021).

84. S. Burow et al., Medaka follicle-stimulating hormone (Fsh) and luteinizing hormone (Lh): Developmental profiles of pituitary protein and gene expression levels. General and Comparative Endocrinology 272, 93–108 (2019).

85. M. R. Royan et al., Functional and developmental heterogeneity of pituitary lactotropes in medaka. General and Comparative Endocrinology 330, 114144 (2023).

86. R. Fontaine, F.-A. Weltzien, Labeling of Blood Vessels in the Teleost Brain and Pituitary Using Cardiac Perfusion with a DiI-fixative. Journal of Visualized Experiments https://dx.doi.org/10.3791/59768, e59768 (2019).

87. F. Wang et al., RNAscope: a novel in situ RNA analysis platform for formalin-fixed, paraffin-embedded tissues. J Mol Diagn 14, 22–29 (2012).

88. M. Golan, L. Hollander-Cohen, B. Levavi-Sivan, Stellate Cell Networks in the Teleost Pituitary. Scientific Reports 6, 24426 (2016).

89. Jamovi (2021) jamovi (Version 2.2.5). in The Jamovi Project (The Jamovi Project, Sydney, Australia).

90. A. Takeuchi, K. Okubo, Post-Proliferative Immature Radial Glial Cells Female-Specifically Express Aromatase in the Medaka Optic Tectum. PLOS ONE 8, e73663 (2013).

